# Bispecific CD20xCD40 Antibodies Achieve Multi-Lineage Modulation of Humoral and Cellular Immunity

**DOI:** 10.1101/2025.11.04.686640

**Authors:** Ben McLaughlin, Mark De Boer, Jeffery Cowden, Modassir Choudhry

## Abstract

Monoclonal antibodies targeting CD20 and CD40 modulate humoral and cellular immunity and have shown therapeutic benefit in B cell malignancies and autoimmune diseases. Here, we describe the generation and preclinical evaluation of two novel bispecific antibodies (DFI201 and DFI205) that combine CD20-mediated B cell depletion inhibition of CD40-mediated influencing populations of B cells, myeloid cells, and T cells. Both bispecifics retained potent B cell cytotoxicity and uniquely extended depletion to non-classical monocytes. A study in cynomolgus macaques showed that administration of bispecific at 5 and 50 mg/kg, or combination therapy with ofatumumab and the anti-CD40 mAb DFI105, achieved rapid and sustained depletion of CD19^+^CD20^+^ B cells in blood and lymphoid tissues, with superior memory B cell clearance compared to monospecific controls Both bispecifics transiently reduced non-classical monocytes and potently depleted dendritic cell subsets (DC1 and DC2). The bispecifics expand regulatory T cells and induce T cell exhaustion without activating T Cells. No cytokine release syndrome or adverse safety signals were observed. Functional immunization with keyhole limpet hemocyanin demonstrated that CD40 engagement was essential to suppress KLH-specific IgG formation. These results establish CD20*×*CD40 bispecific antibodies as a safe and effective approach to achieve deep, multi-lineage immunomodulation with potential for autoimmune indications.

## Introduction

Immune tolerance is a dynamic state in which the immune system is unresponsive to self-antigens, and non-threatening external stimuli while maintaining the ability to defend against pathogens. As established in work earning the 2025 Nobel Prize in medicine, the equilibrium requires coordinated actions of central and peripheral tolerance mediated through specialized populations of immune cell populations such as regulatory T cells, myeloid-derived suppressor cells, and tolerogenic dendritic cells^1^. Rebalancing this system after the emergence of autoimmune disease, or after a transplant, has become a key goal in the development of immunotherapies.

Monoclonal antibodies targeting CD20 and CD40, markers components of the B cell immune system, have emerged as powerful tools in modulating humoral and cellular immune responses, particularly in the context of autoimmune diseases and transplant rejection, as well as important targets in the treatment of B cell lymphomas. These therapies decrease antigen presentation and subsequent T cell activation^2^.

CD20 is a transmembrane cellular protein that has been validated as a therapeutic target expressed by over 95% of B cell lymphocytes throughout their development, from the pre-B cell stage until differentiation into plasma cells, but absent on the hematopoietic stem cell^3^, and present on a small proinflammatory subset of CD8 T cells^4^. CD40 is a membrane-bound costimulatory receptor and a member of the tumor necrosis factor receptor (TNFR) family, bridging innate and adaptive immunity^5^. Like CD20, CD40 is found on lymphocytes, but it is expressed across a wider range of cell types from the pro-B phase through to the plasma cell phase^6–8^. CD40 is also constitutively expressed on dendritic cells. Upon cell activation, the protein is broadly expressed on hematopoietic cells, monocytes and macrophages, and even T cells in cases of autoimmune disease, as well as non-hematopoietic cells, such as endothelial cells (ECs) and CNS resident cells^9,10^.

The CD20 membrane antigen has no physical ligand. Its physiological function is not well understood. CD20 has a role in transmembrane Ca^2+^ conductance^11^ and functions as an organizer of nanoclusters on the B cell membrane^12^. The presence of CD20 is necessary for the function of B cell receptor signaling and activation, but it does not signal independently^13^.

The interaction between CD40 and its ligand CD40L (CD154) is one of the most studied costimulatory signal pathways between B cells, T cells, macrophages and dendritic cells. This pathway has been shown to regulate B cell activation, class switching, antigen presentation, the formation of germinal centers, and humoral memory response^14^. Expression of receptor and ligand has been observed in either cell type; however, the receptor, CD40, is primarily expressed by B cells, and its ligand, CD40L, is common to activated T cells^15^. Stimulation of CD40 initiates, amplifies, and prolong immune responses by suppressing Treg expansion^16^. CD40-CD40L signaling activates both innate and adaptive immune cells, canonically via recruitment of TRAFs and NF-*κ*B activation. Downstream signaling also influences JAK3/STATs, PI3K/AKT, and MAPKs signal pathways^17,18^. The CD40-CD40L dyad also participates in regulating thrombosis, tissue inflammation, hematopoiesis, tumor cell fate, and induction of macrophage effector functions^5,19^. Antagonistic antibody therapies targeting CD40 or its ligand promote expansion and functional maintenance of Tregs by silencing inflammatory costimulatory signals^20^.

Despite the interrelated immunomodulatory mechanisms, Concurrent treatment with anti-CD20 and either agonistic or antagonistic anti-CD40 antibodies has not been extensively explored outside of some isolated lymphoma ^21,22^ and xenotransplantation studies^23^. Activation of CD40 has been shown to downregulate CD20 expression^24^. The opportunity for synergistic benefit of concurrent treatment, or treatment with a dual-targeting anti-CD20/CD40 bispecific, is significant.

We investigate the profound and multifaceted effects on cellular and humoral adaptive immunity associated with concurrent B cell depletion and minimizing myeloid cell activation and engagement by inhibiting CD40. Our test articles include using monospecific antibodies targeting CD20, a CD40 antagonist DFI105, and DFI205, a novel bispecific antibody combining effector-mediated depletion of B cells with inhibition of CD40.

## Methods

### Novel Antibodies

Novel antibody test articles include DFI105, an IgG4 antagonistic anti-CD40 mAb that allosterically inhibits CD40. This antibody shares a binding epitope with its precursor mAbs ch5D12, and PG102 and fully retains its unique mode of action^19,25^. This CD40 binding epitope is conserved in humans, marmosets, Rhesus, and Cynomolgus monkeys^26^. The two novel bispecific antibodies combine the anti-CD40 binding of the epitope of DFI105 with anti-CD20 targeting domains and Fcs selected to optimize for ADCC (DFI201) and CDC (DFI205).

### DFI105

DFI105 is an allosteric CD40 antagonistic deimmunized monoclonal antibody. Several anti-body engineering steps were undertaken to optimize the original 5D12 mouse monoclonal antibody (ref) for human use. These steps included deimmunization, optimizing expression levels in CHO cells, introducing the VHI29L back mutation, and removal of deamidation aggregation liabilities.

DFI105 binds to an epitope on non-human CD40 on an outward-facing loop, positioned away from the known CD40-CD40L interaction site^27^. It can completely inhibit the CD40L-CD40 signal without competing with CD40L for receptor occupancy^25^. The molecule provides an allosteric inhibition of C40-CD40L signaling. Full activation of this pathway requires force-modulated mechanotransduction of the catch-slip bond between CD40 and CD40L, as well as conformational changes to CD40 forming higher-order structures^27–29^.

### PBMC Killing Assay

Antibodies were serially diluted (40 *µ*g/mL start, seven five-fold steps) in complete medium and added to PBMCs (final volume 200 *µ*L). After 24 h incubation at 37 *^◦^*C, cells were harvested for staining.

Cells were washed, stained with Live/Dead dye, blocked with Fc Block, then labeled with fluorochrome-conjugated antibodies against CD45, CD19, CD20, and other lineage markers. Count-Bright beads were added before acquisition on a BD Fortessa cytometer.

Live singlet CD45^+^ lymphocytes were gated; B cells were CD19^+^CD20^+^. Absolute counts were calculated via bead ratios, and percent depletion versus no-antibody control was plotted in GraphPad Prism to derive EC_50_ values. Assays were run in triplicate with at least three donors.

### Animals

Cynomolgus macaques (Macaca fascicularis), 2.5–4 kg in weight, comprising equal numbers of males and females (total n=18), were used. All animals were naïve to previous treatments and underwent a 14-day acclimatization period under controlled conditions (20–26◦C, relative humidity 30–70%, 12-hour light/dark cycle). Animals were individually housed during the Experimental phase and provided with unrestricted access to food (standard monkey diet) and filtered municipal drinking water. All animal procedures were approved by the Institutional Animal Care and Use Committee (IACUC) at PharmaLegacy Laboratories and conducted in compliance with ethical standards.

### Primate Study Experimental Design

Animals were randomly allocated to nine groups (n=2 per group; one male and one female), balanced by body weight. Treatments involved intravenous administration via brachiocephalic or posterior saphenous veins of novel bispecific antibodies DFI207 or DFI205 at doses of 5 or 50 mg/kg, monospecific antibodies (Rituxan, Kesimpta, Anti-CD40) at 5 mg/kg, a combination therapy (Kesimpta+Anti-CD40), or vehicle control (PBS). Treatments were administered on Day 0 and Day 7. Animals were immunized with keyhole limpet hemocyanin (KLH, 10 mg i.m.) on Day 8 for functional immunological assessment.

### Sampling And Assays

Blood samples were collected at defined intervals (Days -3, 0 pre-dose, 1, 7, 14, 21) for hematology, clinical chemistry, pharmacokinetic (PK), pharmacodynamic (PD), and anti-drug antibody (ADA) analyses. Additional sampling included KLH-specific antibody quantification and complement activation assessments at selected timepoints. Bone marrow, lymph nodes, and spleen tissues were harvested on Days 14 and 21 for flow cytometry and immunohistochemistry (IHC).

Flow cytometry characterized B-cell subsets, T/NK subsets, dendritic cells, and monocytes in blood and tissues. PD parameters assessed included cytokines (IL-1*β*, IL-2, IL-6, IL-10, IL-12p70, IFN-*γ*, TNF-*α*, G-CSF, IL-8), immunoglobulin isotypes (IgM, IgG, IgA), and inflammatory mediators (IFN*β*, MMP3, BLC, CD30, sCD40). KLH-specific IgG and IgM responses were measured by ELISA. PK bioanalysis utilized a validated generic human IgG Fc ELISA, with non-compartmental parameters calculated using WinNonlin software.

### Flow Cytometry Analysis

Flow cytometry characterized immune cell populations including B-cell subsets (activated B cells, memory B cells, mature naïve B cells, plasma cells), T/NK subsets (Tregs, exhausted T cells, naïve and effector T cells), dendritic cells (DC1, DC2), and monocytes (classical and non-classical) in peripheral blood, bone marrow, lymph nodes, and spleen. Samples were stained using multi-color antibody panels targeting markers such as CD3, CD4, CD8, CD19, CD20, CD25, CD27, CD45, CD56, CD69, PD-1, CCR7, and HLA-DR. Data acquisition was performed on CytoFlex LX and BD Fortessa flow cytometers, followed by analysis using FlowJo software.

The gating strategy involved an initial gate based on forward scatter (FSC) and side scatter (SSC) to identify lymphocyte populations. Singlet cells were gated using FSC-A versus FSC-H plots. Live cells were identified through the exclusion of viability dye-positive cells.

Panel 1 T cells:

- Naïve T cells (CD3+, CD4+/CD8+, CD45RA+, CCR7+)
- Central Memory T cells (CD3+, CD4+/CD8+, CD45RA-, CCR7+, CD62L+)
- Effector Memory T cells (CD3+, CD4+/CD8+, CD45RA-, CCR7-, CD62L-)
- T memory stem cells (CD3+ CD8+ CD45RA+ CCR7+ CD62L+ CD279-)
- TEMRA (CD3+, CD4+/CD8+, CD45RA+, CCR7-)
- Potential Regulatory T cells (CD3+, CD4+, CD25+, CD127low)
- Regulatory T cells (CD3+, CD4+, CD25+, PD-1+)
- Exhausted T cells (CD3+, PD-1+)
- T helper :(CD3+, CD4+ CD69+ MFI)
- T killer: (CD3+, CD8+ CD69+ MFI)
- Activated T Cells (CD3+, CD25+)
- CD56+ T cells (CD3+, CD56+)
- NK Cells (CD3-CD56+)

Panel 2: B cells:

- Pro-B (CD19+, CD34+, CD10+, CD38hi)
- Pre-B (CD19+, CD34-, CD10+, CD38hi)
- transitional B cells (CD10+, CD38+, IgM*+)*
- T1 (CD19+, CD38hi, CD10+, CD21low)
- T2 (CD19+, CD38hi, CD10+, CD21+)
- Naïve B cells (CD19+, CD20+, IgM+, CD27-) CD38low
- Unswitched memory B cells (CD19+, CD20+, CD27+)
- Marginal Zone B cells (CD19+, CD20+, CD27+, IgM+, CD21hi)
- Mature naive B cells (CD19+, CD20+, CD21hi, CD27-)
- Switched memory B cells (CD19+, CD20+, IgM-, CD27+)
- Exhausted B cells (CD19+, CD21low, CD27-, CD10-)
- Plasmablasts (CD19+, CD20low/-, CD27hi, CD38hi)
- Plasma cells (CD19+/-, CD20-, CD138+, CD38hi)
- Marginal zone B cells (CD19+, CD20+, IgM+, CD21+, CD23-)
- Follicular B cells (CD19+, CD20+, CD21+, CD23+)
- Activated B cells (CD19+, CD20+, CD38+)
- Regulatory B cells (CD19+, CD24hi, CD38hi)
- B1 cells (CD19+, CD20+, CD43+, CD27+)

Panel 3: Myeloid Cells and CD20+ T Cells

Monocyte Subsets

- Classical Monocytes (CD14++, CD16-)
- Intermediate Monocytes (CD14++, CD16+)
- Non-classical Monocytes (CD14+, CD16++)
- Dendritic Cell Populations
- Conventional DC1 (HLA-DR+ & MFI, CD14-, CD11c+, CD123-)
- Conventional DC2 (HLA-DR+ & MFI, CD14-, CD11c+, CD123low)
- Plasmacytoid DC (HLA-DR+ & MFI, CD14-, CD11c-, CD123

T Cells

- CD3+CD20+ T cells

### Anti-Drug Antibody (ADA) Assay

A bridging assay detected ADAs. Serum samples collected on Days 0, 7, 14, and 21 were initially screened for the presence of antibodies capable of bridging biotinylated and SULFO-TAG labeled therapeutic proteins. Samples surpassing a predetermined cut-point were considered putatively positive. Confirmation assays involved an antigen-specific monkey IgG ELISA to confirm the presence of specific immunoglobulin responses against therapeutic constructs.

### Pharmacodynamics and Pharmacokinetics

PD parameters assessed included cytokines (IL-1*β*, IL-2, IL-6, IL-10, IL-12p70, IFN-*γ*, TNF-*α*, G-CSF, IL-8), immunoglobulin isotypes (IgM, IgG, IgA), and inflammatory mediators (IFN*β*, MMP3, BLC, CD30, sCD40). KLH-specific IgG and IgM responses were measured by ELISA. PK bioanalysis utilized a validated generic human IgG Fc ELISA, with non-compartmental parameters calculated using WinNonlin software

### Safety and Tolerability

Safety parameters, including body weight, body temperature, clinical observations, complete blood counts (CBC), clinical chemistry, and coagulation parameters, were monitored regularly. Animals showing adverse symptoms or significant weight loss were immediately evaluated.

### Statistical Analysis

Data were expressed as mean *±* SEM. Statistical comparisons utilized two-way ANOVA followed by Dunnett’s test, with significance set at p<0.05.

## Results

### In Vitro Cell Killing

The novel bispecific anti-CD20/CD40 antibodies combined the B cell depletion characteristics of CD20 antibodies with CD40. The bispecificity did not result in a significant reduction of B cell depletion EC50, and expanded the depletable cell types to include non-classical monocytes. The antibody function was conserved across human and cyno PBMCs, Figures 1 and 2, Tables 1 and 2.

**Figure 1:**
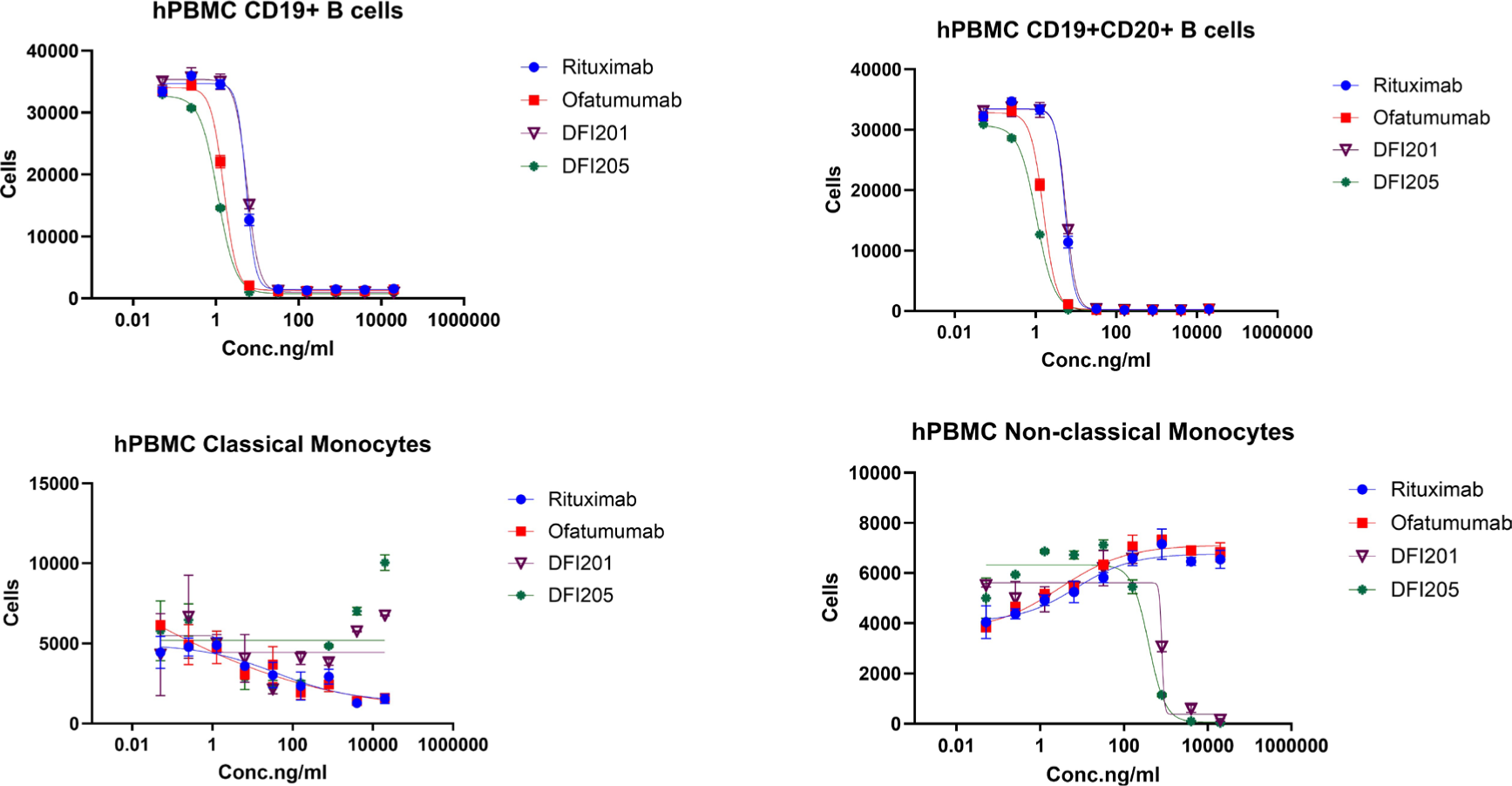
In Vitro Cell Killing of hPBMCs EC50s were.

**Figure 2:**
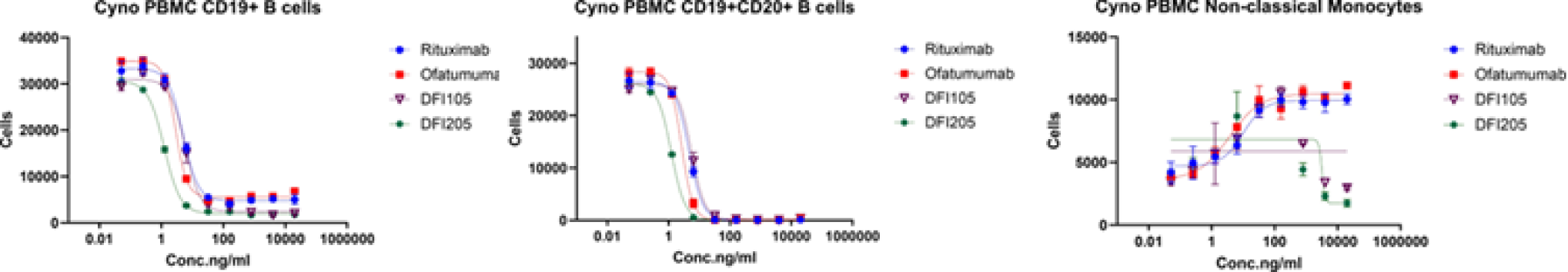
In Vitro Cell Killing in Cynomolgus Monkey PBMC.

**Table 1:**
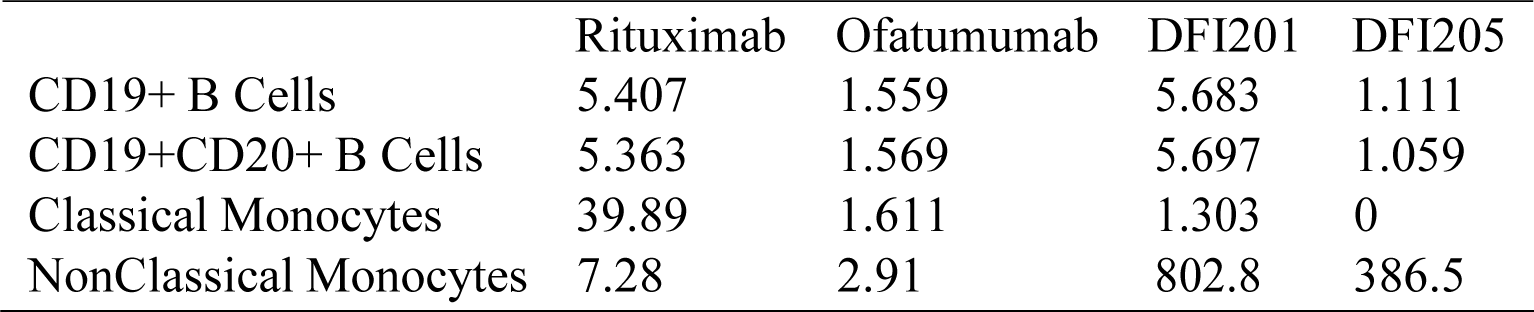
EC50 of Human Cell Killing.

**Table 2:**
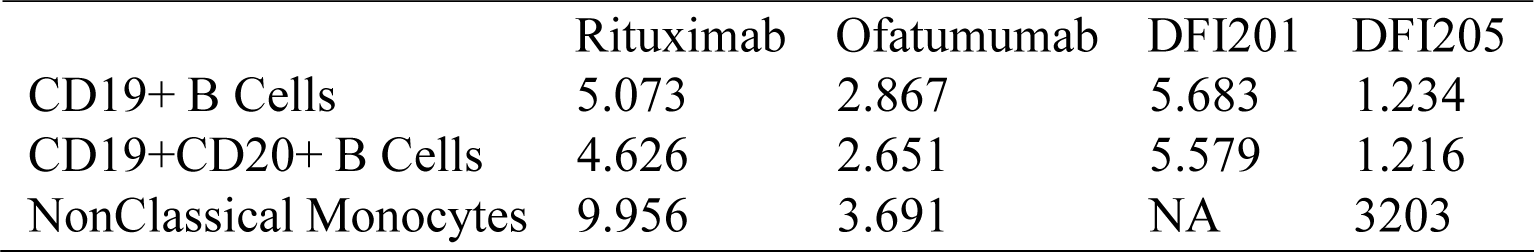
EC50 of PBMC Cell Killing.

### Pharmacokinetics

The serum concentration-time profiles of bispecific antibodies along with monospecific controls showed dose-dependent pharmacokinetics, Table 3. Higher doses demonstrated prolonged serum exposure and greater area under the curve (AUC). The bispecifics showed shorter half-lives than either anti-CD20 or anti-CD40 monospecific antibodies.

**Table 3:**
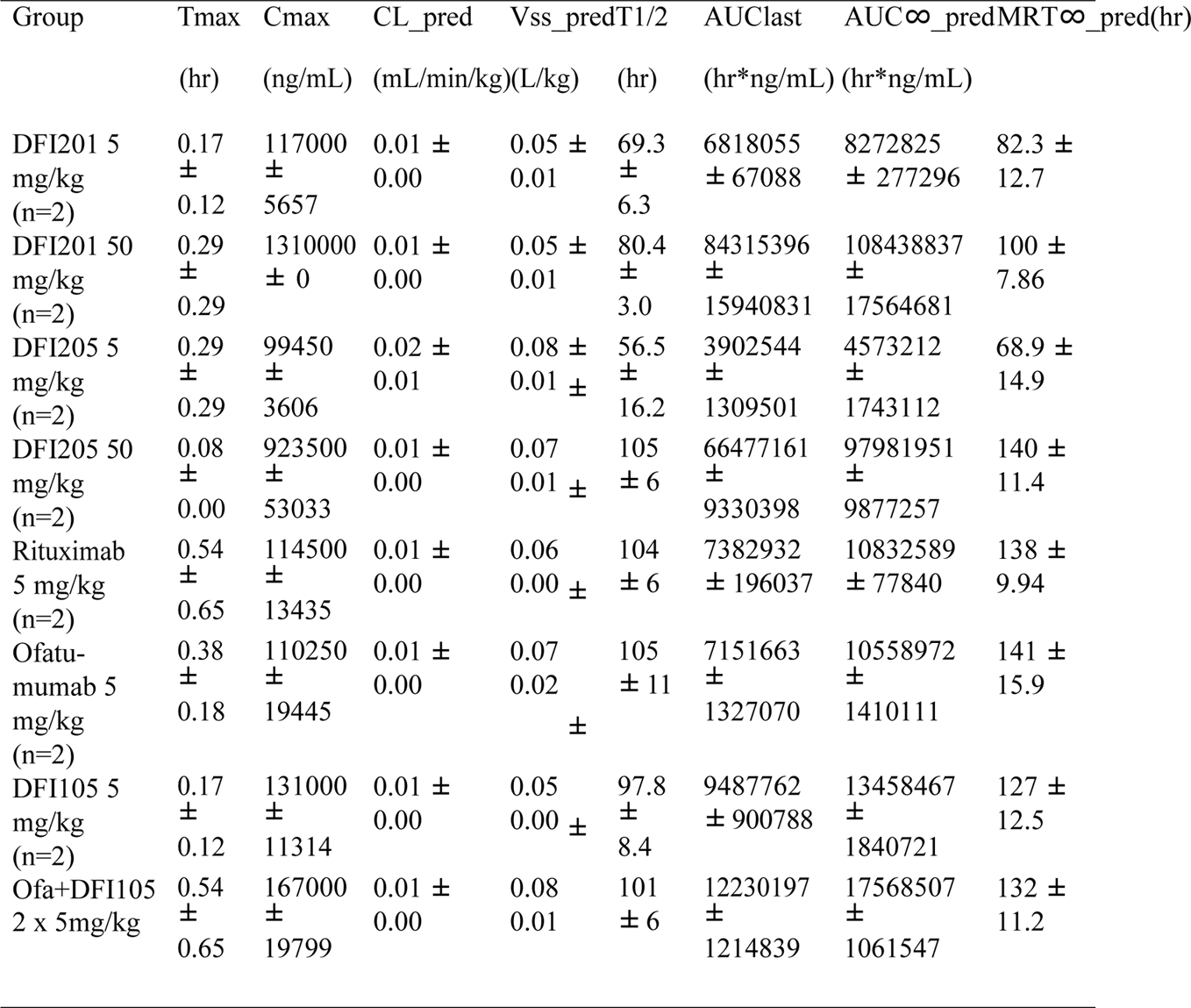
Pharmokinetic Parameters of anti-CD20, and anti-CD40 antibodies and bispecifics.

| Group | Tmax | Cmax | CL_pred | Vss_pred | T1/2 | AUClast | AUC∞_pred | MRT∞_pred(hr) |
| --- | --- | --- | --- | --- | --- | --- | --- | --- |
|  | (hr) | (ng/mL) | (mL/min/kg) | (L/kg) | (hr) | (hr*ng/mL) | (hr*ng/mL) |  |
| DFI201 5 mg/kg (n=2) | 0.17 ± 0.12 | 117000 ± 5657 | 0.01 ± 0.00 | 0.05 ± 0.01 | 69.3 ± 6.3 | 6818055 ± 67088 | 8272825 ± 277296 | 82.3 ± 12.7 |
| DFI201 50 mg/kg (n=2) | 0.29 ± 0.29 | 1310000 ± 0 | 0.01 ± 0.00 | 0.05 ± 0.01 | 80.4 ± 3.0 | 84315396 ± 15940831 | 108438837 ± 17564681 | 100 ± 7.86 |
| DFI205 5 mg/kg (n=2) | 0.29 ± 0.29 | 99450 ± 3606 | 0.02 ± 0.01 | 0.08 ± 0.01 | 56.5 ± 16.2 | 3902544 ± 1309501 | 4573212 ± 1743112 | 68.9 ± 14.9 |
| DFI205 50 mg/kg (n=2) | 0.08 ± 0.00 | 923500 ± 53033 | 0.01 ± 0.00 | 0.07 ± 0.01 | 105 ± 6 | 66477161 ± 9330398 | 97981951 ± 9877257 | 140 ± 11.4 |
| Rituximab 5 mg/kg (n=2) | 0.54 ± 0.65 | 114500 ± 13435 | 0.01 ± 0.00 | 0.06 ± 0.00 | 104 ± 6 | 7382932 ± 196037 | 10832589 ± 77840 | 138 ± 9.94 |
| Ofatu-mumab 5 mg/kg (n=2) | 0.38 ± 0.18 | 110250 ± 19445 | 0.01 ± 0.00 | 0.07 ± 0.02 | 105 ± 11 | 7151663 ± 1327070 | 10558972 ± 1410111 | 141 ± 15.9 |
| DFI105 5 mg/kg (n=2) | 0.17 ± 0.12 | 131000 ± 11314 | 0.01 ± 0.00 | 0.05 ± 0.00 | 97.8 ± 8.4 | 9487762 ± 900788 | 13458467 ± 1840721 | 127 ± 12.5 |
| Ofa+DFI105 2 x 5mg/kg | 0.54 ± 0.65 | 167000 ± 19799 | 0.01 ± 0.00 | 0.08 ± 0.01 | 101 ± 6 | 12230197 ± 1214839 | 17568507 ± 1061547 | 132 ± 11.2 |

### Pharmacodynamics Cytokine Release

No evidence of cytokine release syndrome was observed in any treatment group.

Monkeys treated in the study were evaluated for evidence of cytokine release syndrome on day -3, 0, 1hr, 7, 14, and 21. Levels of IFN-*γ*, G-CSF, TNF-*α*, IL-1*β*, IL-2, IL-6, IL-10, and IL12p70 remained consistent with both baseline and vehicle controls in all groups. One monkey treated with the combination of Ofatumumab and DFI105 showed an increase in IL-8 on Days 7 and 21 to 30 pg/mL. These levels are not believed to be related to treatment, as the other monkey in the dose group showed ∼10-fold lower levels.

### Compliment

Compliment cascade-related proteins were quantified 1 hour after the first dose, and at the termination of the study, Figure 3. The levels of C3a, Factor1, and C5a in serum were all below the detection limit: 36 ng/ml, 2109 ng/ml, and 0.9 ng/ml, respectively. Treatment groups did not differentiate from the vehicle in serum Ba, C4a, and SC5b9. Animals treated with 50 mg/kg bispecific, or 5 mg/kg of Ofatumumab had higher Bb than the vehicle group one hour after dosing. There was no differentiation in other groups or at different timepoints. Both bispecific groups and the DFI105 group showed elevated Factor H in serum on day 21, relative to the vehicle.

**Figure 3:**
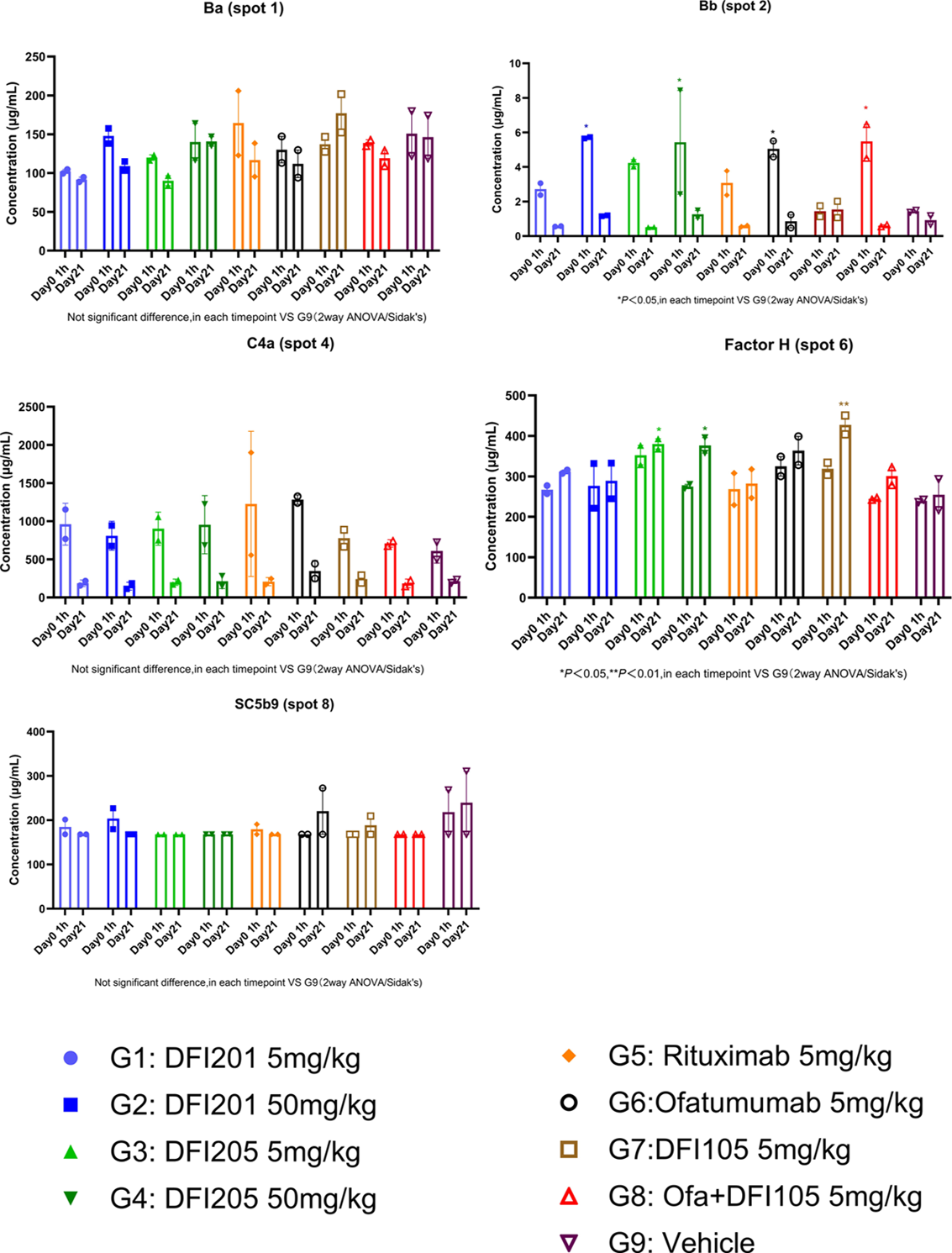
Measurement of compliment in serum at 1h, and on day 21. The level of C3a, Factor1, C5a in serum were all below the detection limit: 36 ng/ml, 2109 ng/ml, 0.9 ng/ml respectively.

### Isotype MSD Analysis

The levels of IgA, IgG, and IgM in serum did not show significant differences when compared with G9 (vehicle) on day 3, day 0, day 1, day 7, day 14, or day 21. No significant change from the pre-treatment baseline was observed, Figures 4, 5, and 6.

**Figure 4:**
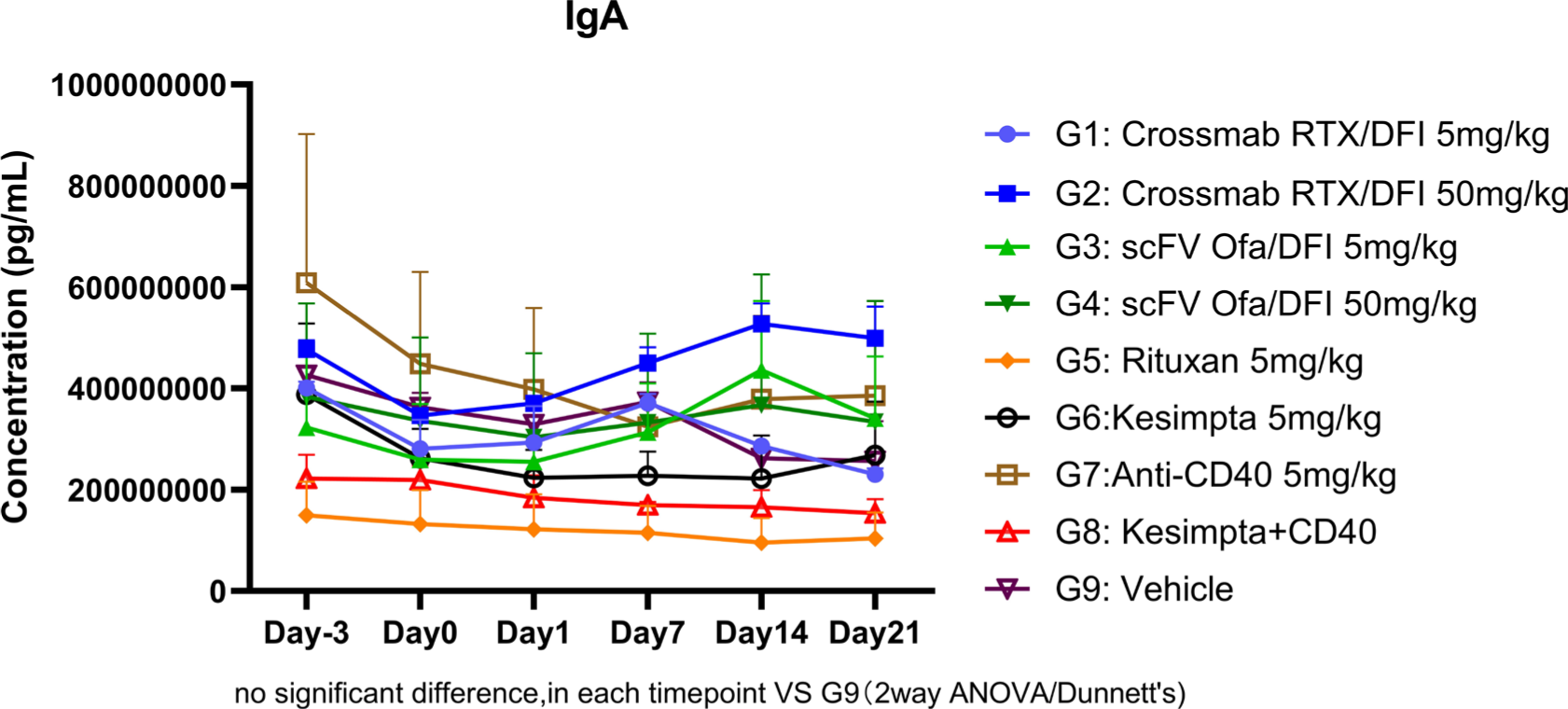
IgA levels in serum during NHP Study.

**Figure 5:**
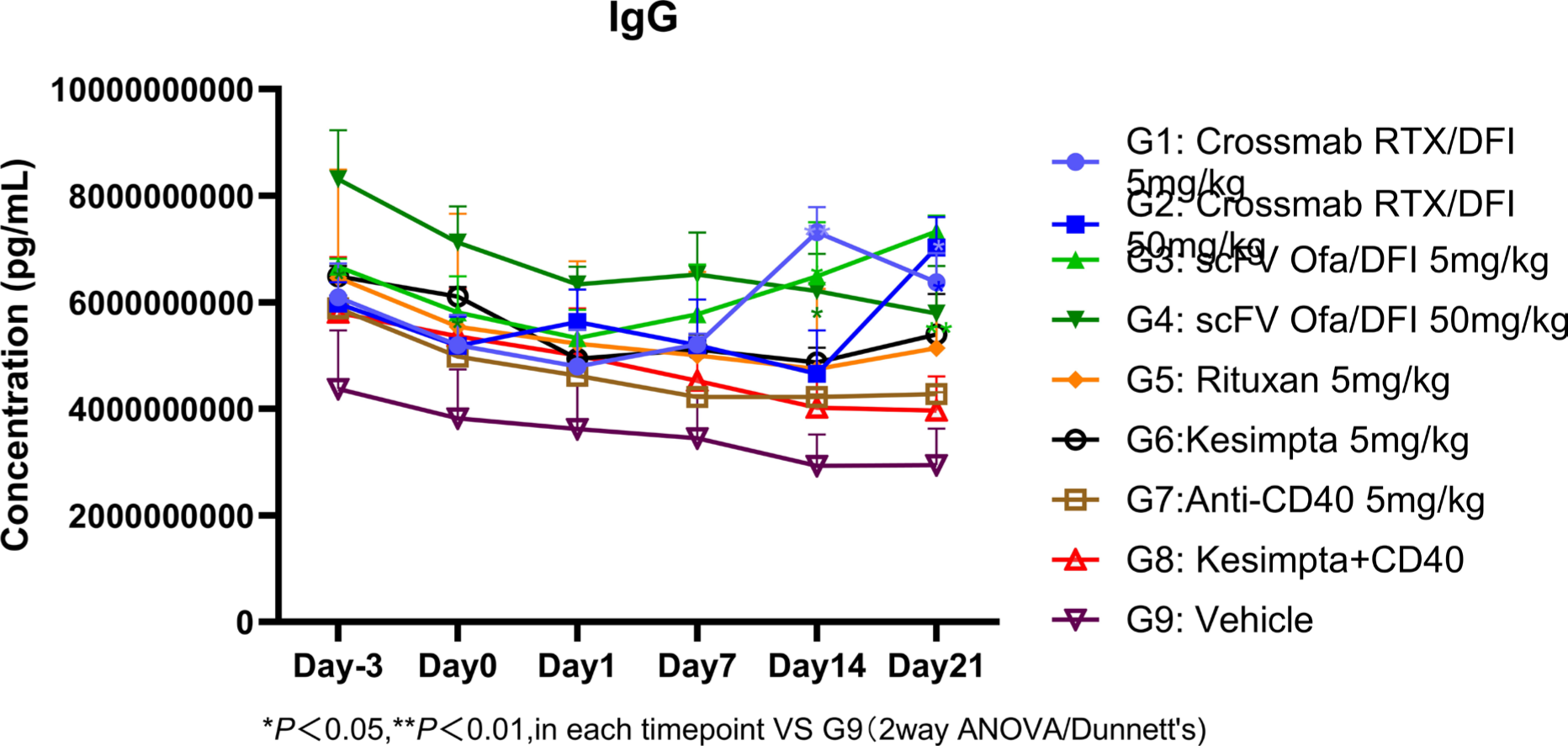
IgG levels in serum during NHP Study.

**Figure 6:**
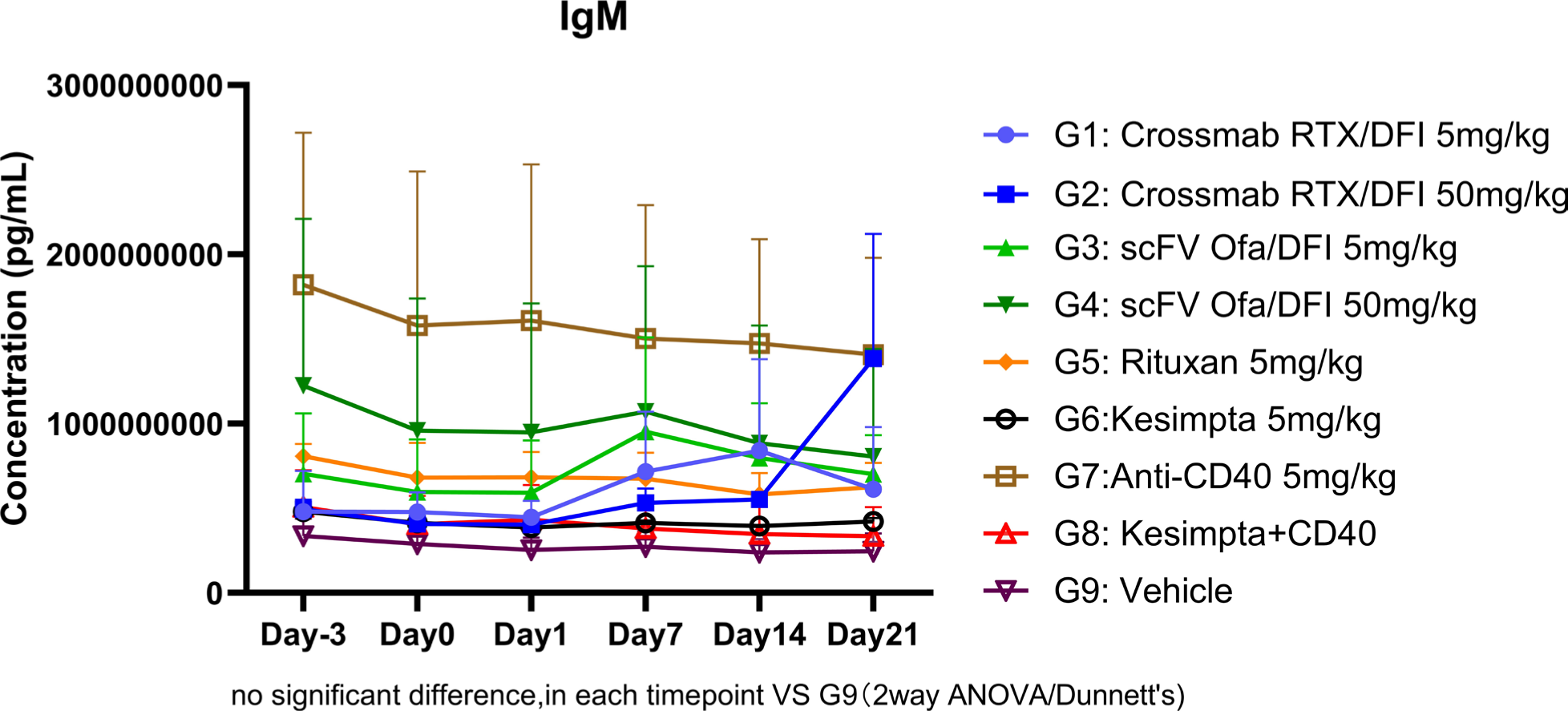
IgM levels in serum during NHP Study.

### KLH Specific IgG Levels in Serum

KLH-specific IgG and IgM were monitored following the abbreviated KLH protocol, Figures 7 and 8. B-cell depleting antibodies alone were not able to inhibit the formation of KLH-specific IgG. This response was only seen after treatment with an anti-CD40, CD20/CD40 bispecific, or combination.

**Figure 7:**
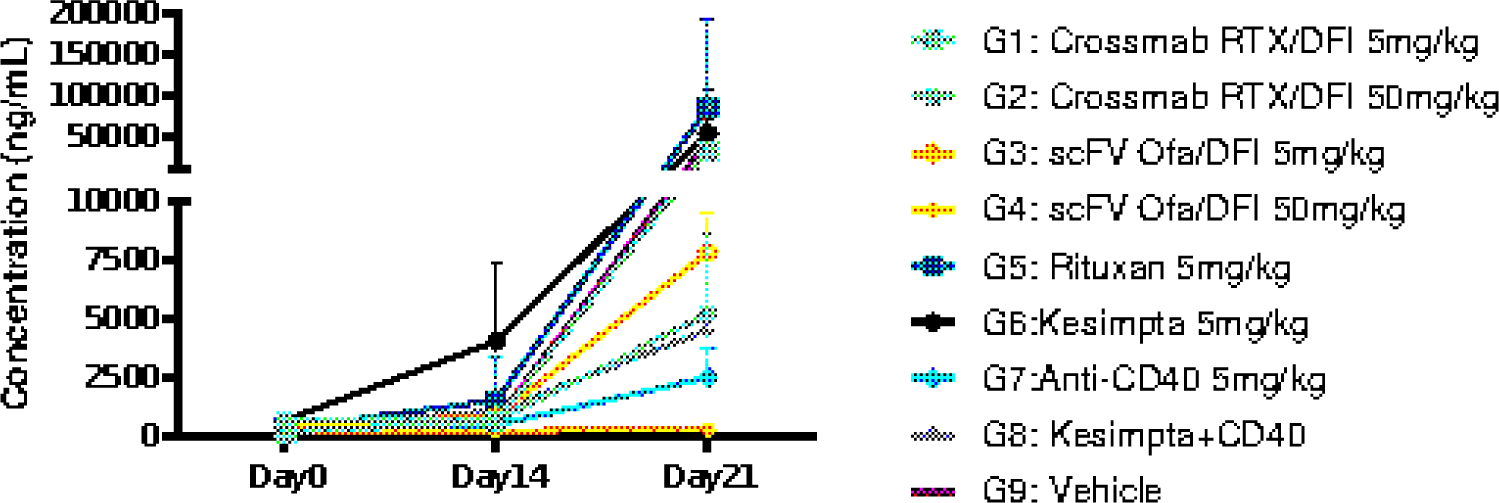
KLH Specific IgG Levels in Serum following a single immunization on Day 8.

**Figure 8:**
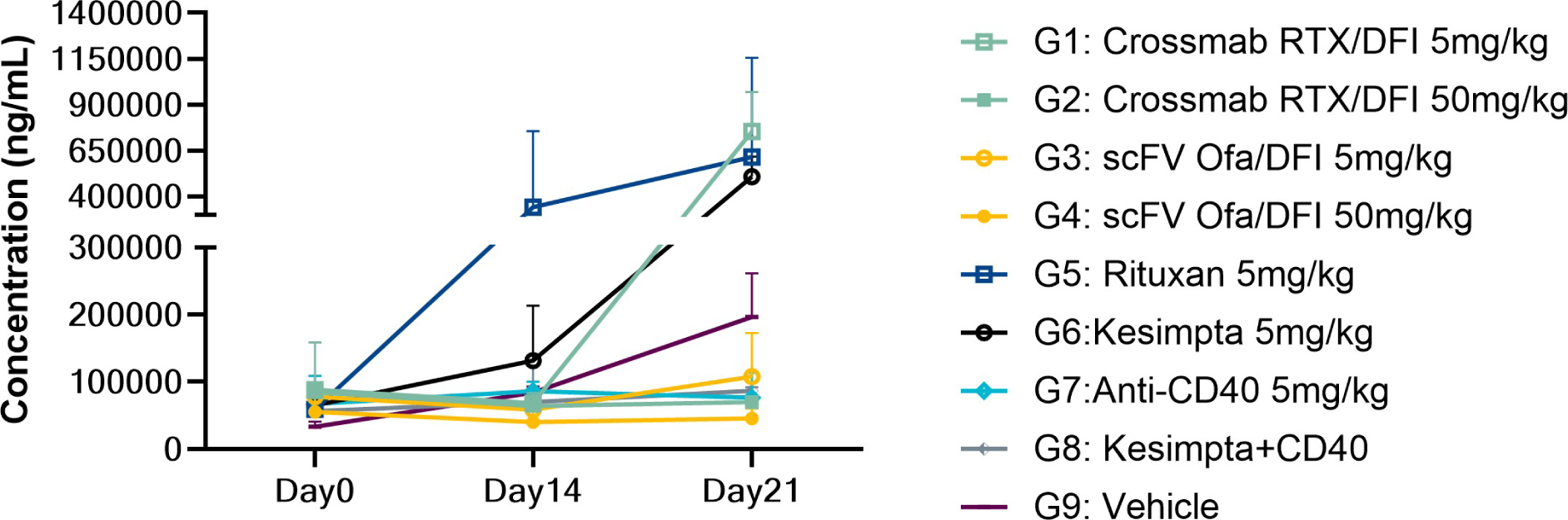
KLH Specific IgM in Serum following a single immunization on Day 8.

#### Flow Cytometry Analysis

Flow cytometry analysis revealed changes in cell types and subtypes during the study. DFI105 was IgG4 and not expected to show any Fc-mediated cytotoxicity. The other Fc active antibodies showed immediate and sustained depletion of circulating B Cells,

Figure 9. Repopulation as observed on day 21 showed some dose dependency, but was shown for all drugs. Only the 50 mg/kg dose of DFI205 and the combination of ofatumumab and DFI105 demonstrated sustained and comprehensive depletion across compartments, Figure 9. Depletion of tissue resident memory B cells by bispecific and combination therapy exceeded the depletion by conventional anti-CD20 mAbs in bone marrow, lymph node, and spleen, Figure 10

**Figure 9:**
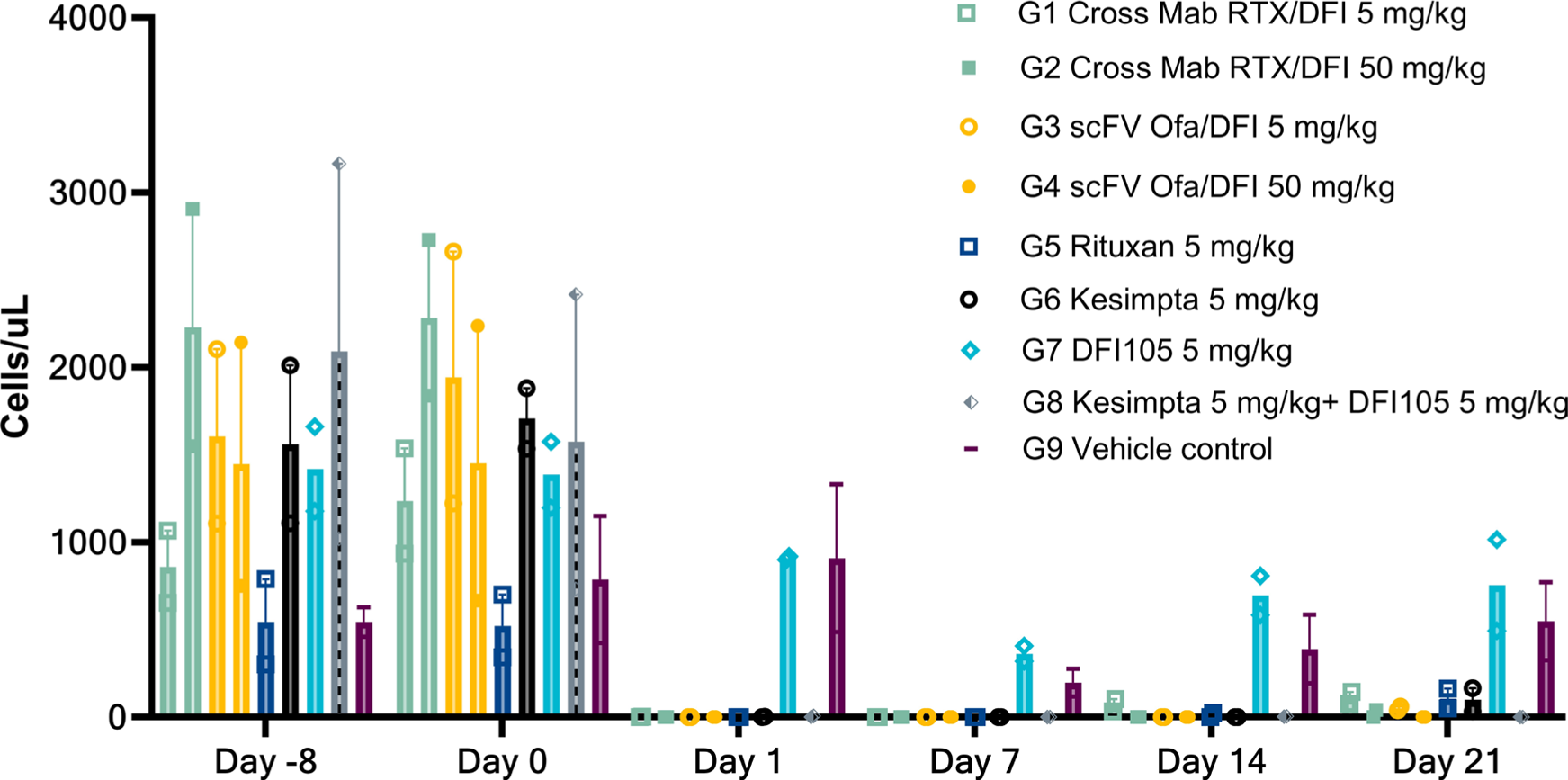
CD19+CD20+ B Cells in the Peripheral Blood.

**Figure 10:**
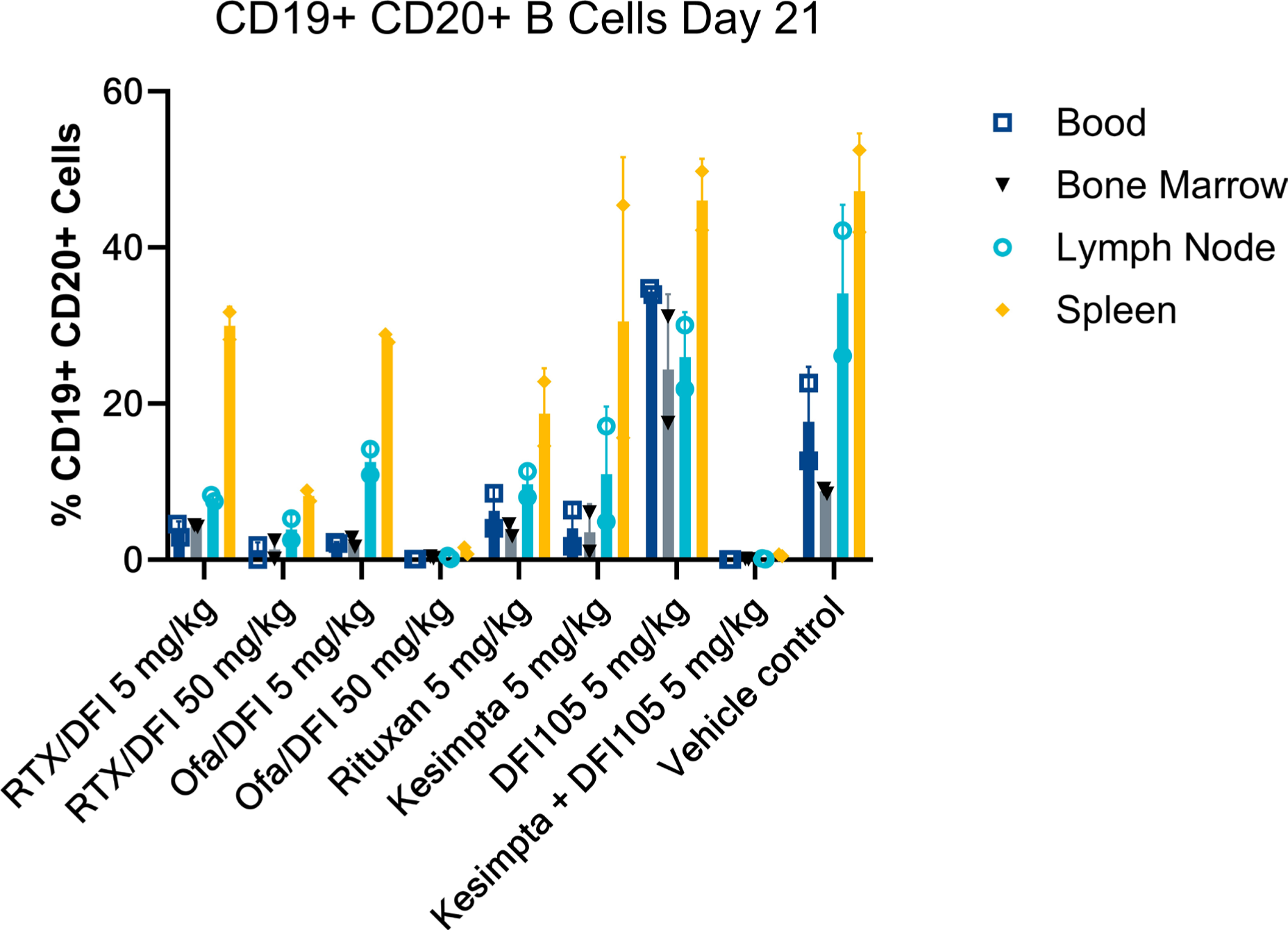
B Cell Populations in in Blood, Bone Marrow, Lymph node and Spleen two weeks after treatment.

**Figure 11:**
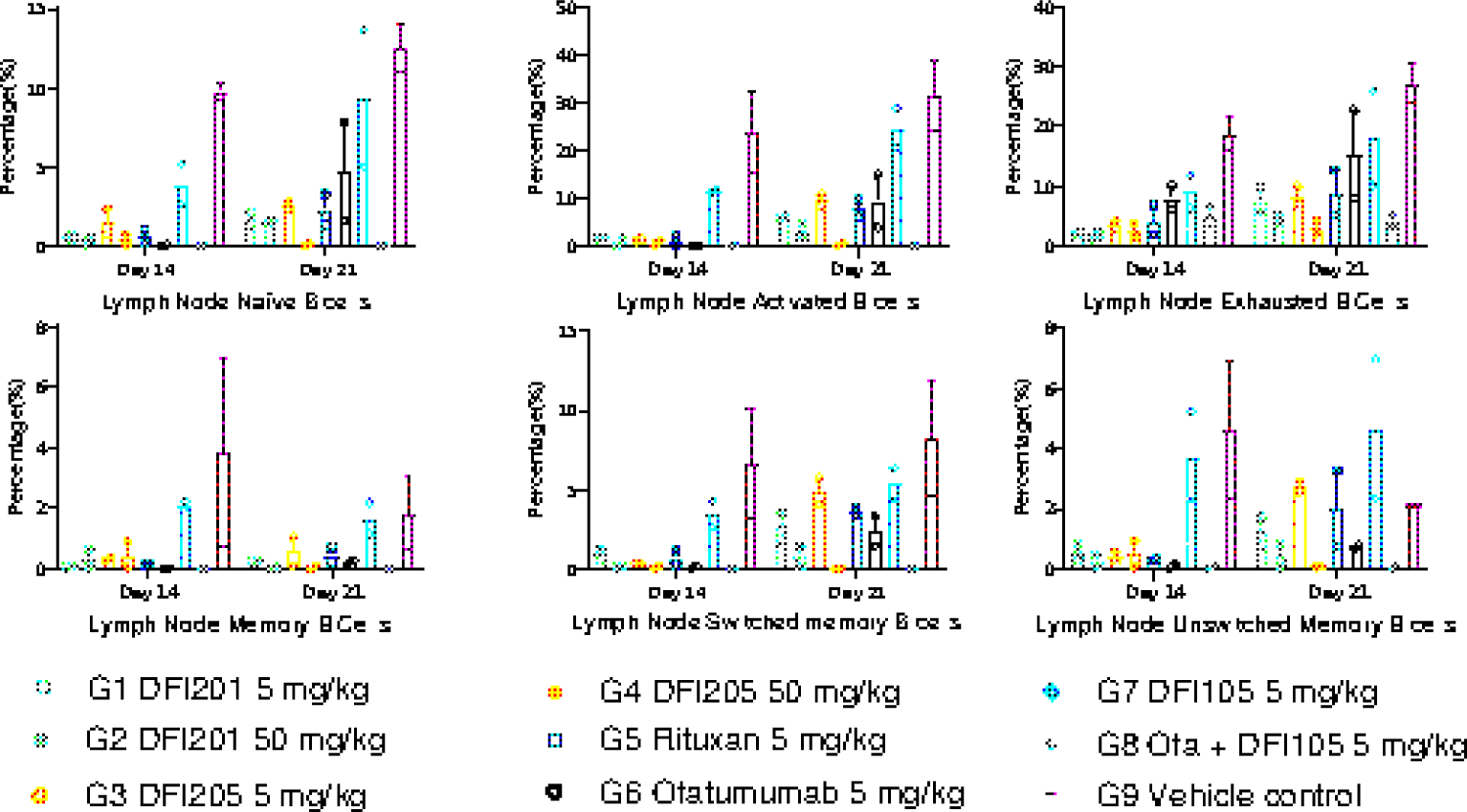
B Cell Sub-Types in the Lymph Nodes One and Two Weeks After Dosing.

**Figure 12:**
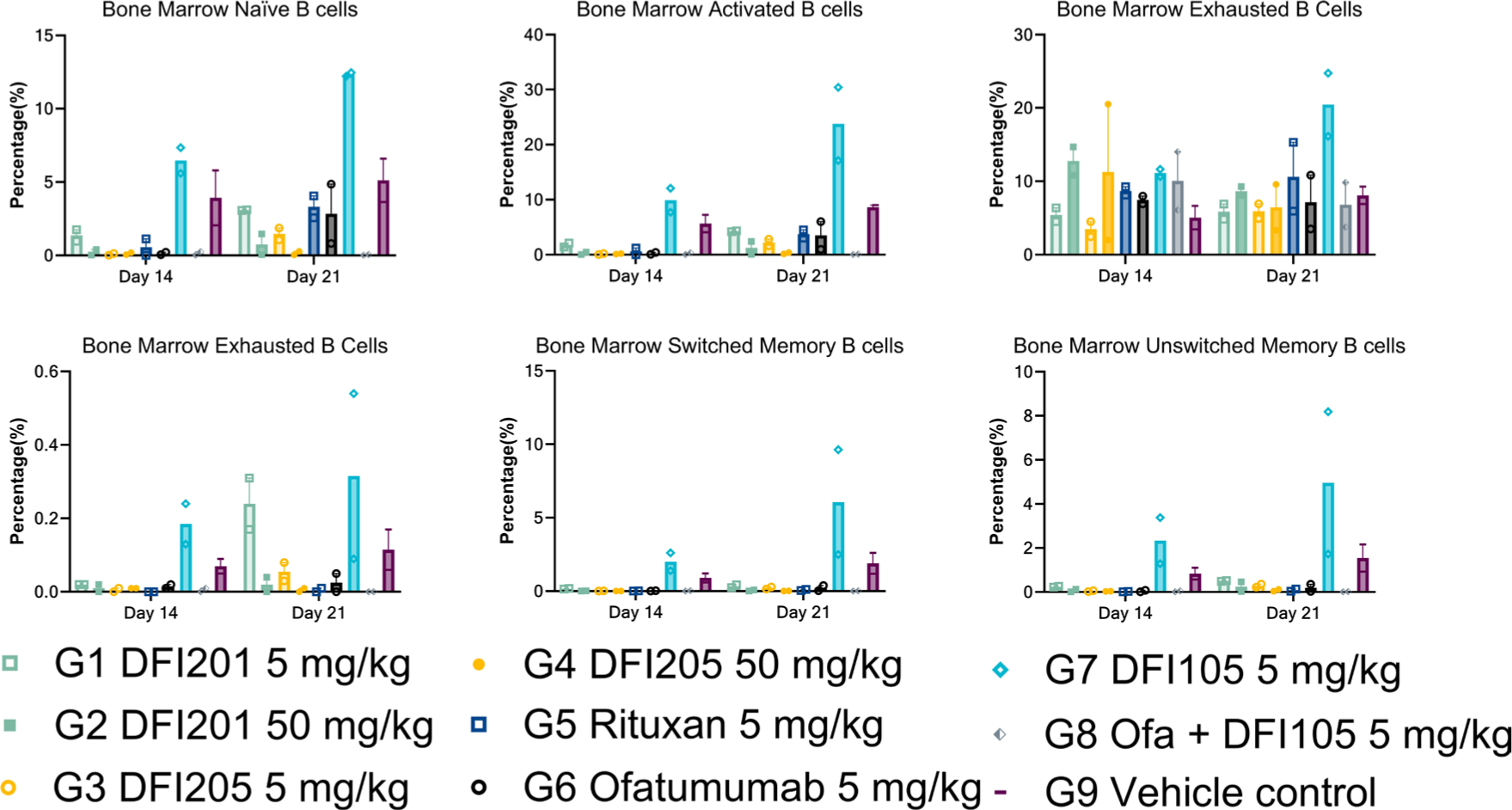
B Cell Sub-Types in the Bone Marrow One and Two Weeks After Dosing.

**Figure 13:**
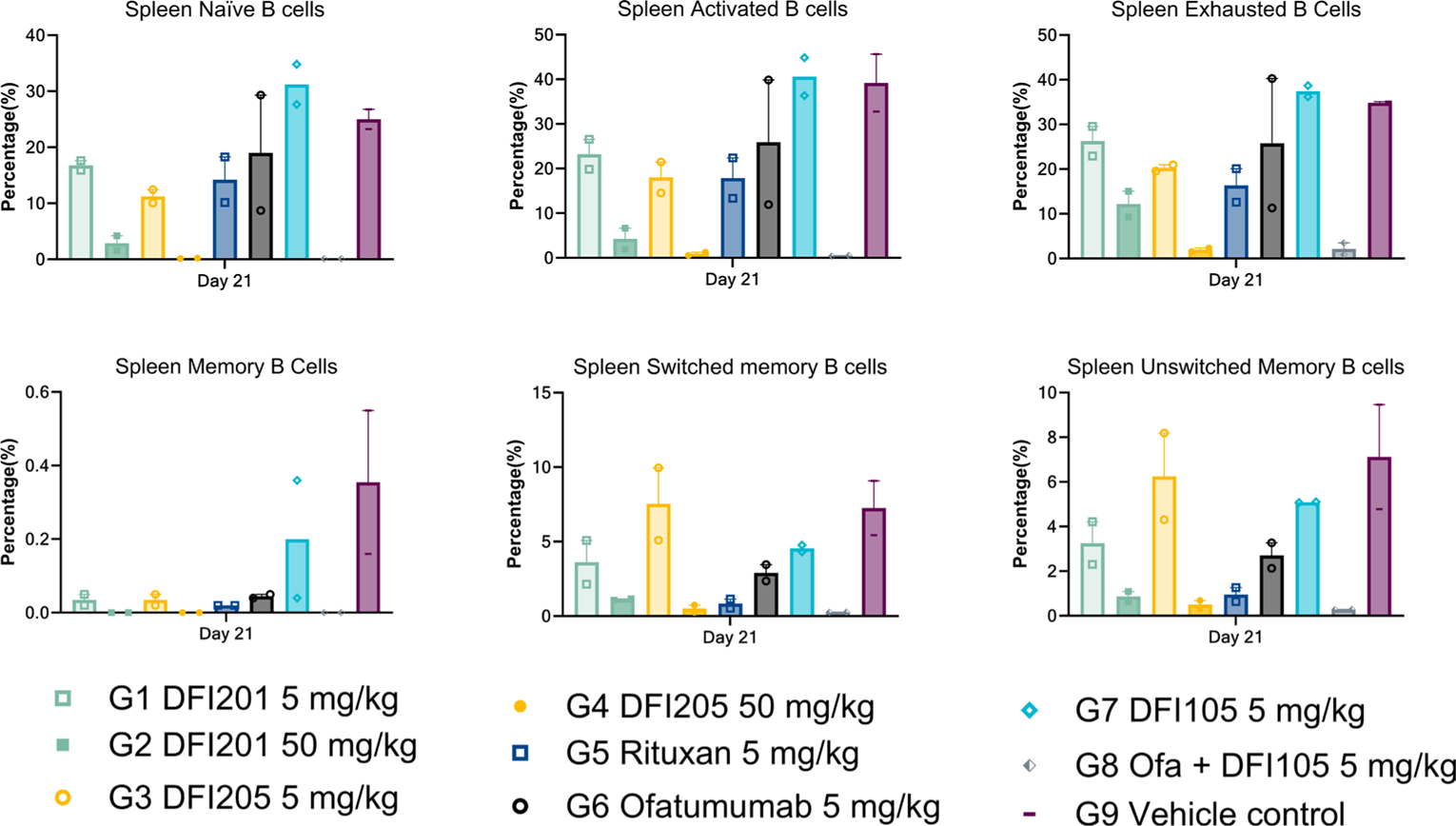
B Cell Sub-Types in the Spleen One and Two Weeks After Dosing.

The population of classical, non-classical, and intermediate monocytes was trended for the duration of the study, Figure 14. The first round of dosing with bispecifics showed an initial decrease in non-classical monocytes before homeostasis returned. The CD20+CD40 bispecifics can deplete non-classical monocytes not killed by either the CD20, CD40, or combination therapy.

**Figure 14:**
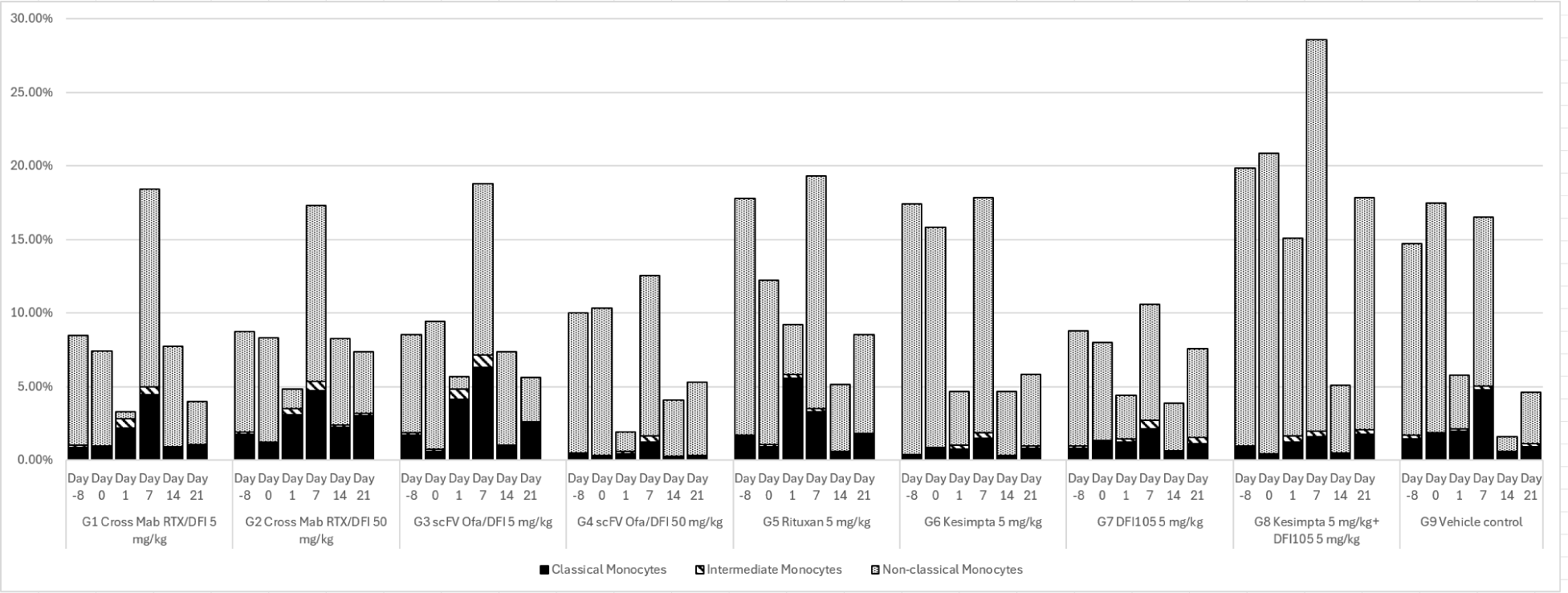
Monocyte Populations During the NHP Study.

Conventional dendritic cell depletion was observed with the bispecific and the anti-CD20s, but not the CD40 or combination treatment, as shown in Figure 15. The anti-CD20s only depleted DC1s, while the BsAbs potently depleted both DC1 and DC2 subsets, Figure 16.

**Figure 15:**
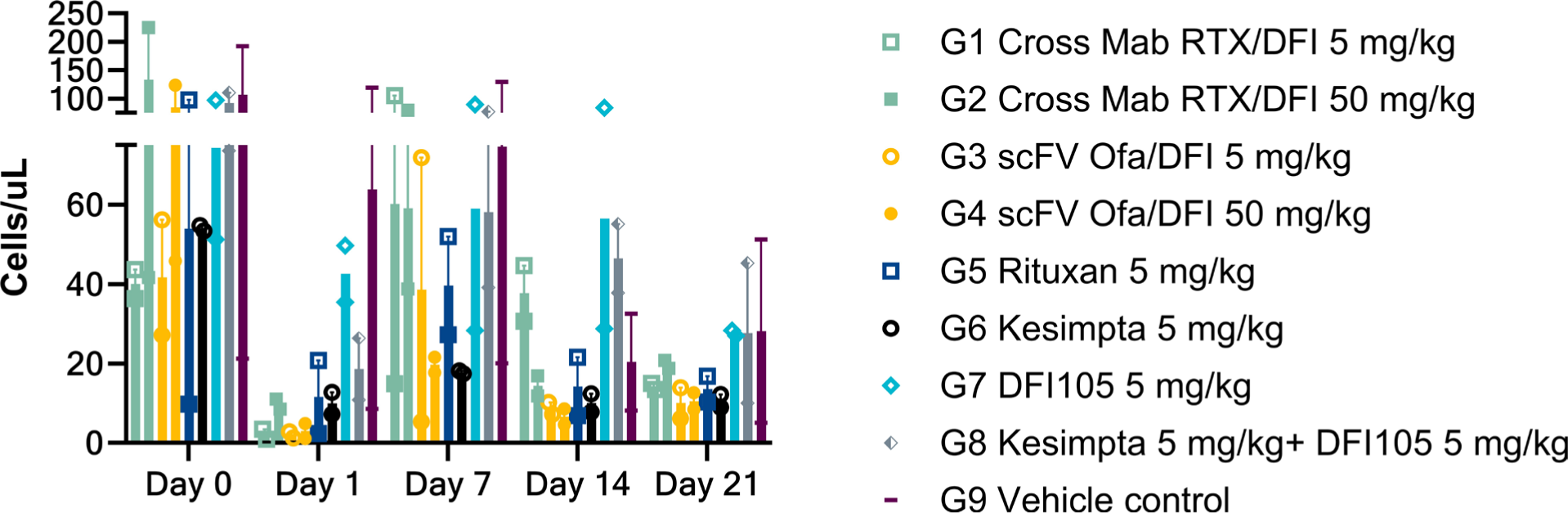
Blood Conventional Dendritic Cell (DC1) Population During the Study.

**Figure 16:**
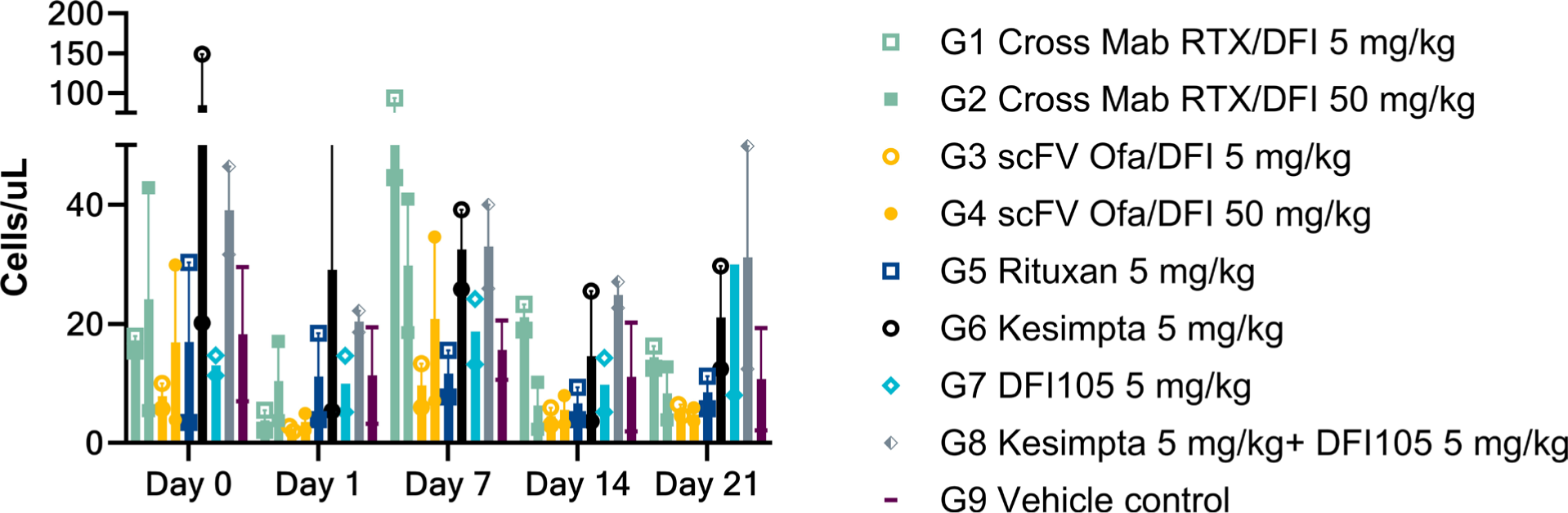
Blood Conventional Dendritic Cell 2 (DC2) Population During the Study.

The relative abundance and distribution of T cell subtypes monitored during the NHP study indicate the immune system modulation induced by the anti-CD20/CD40 therapies. The therapy was shown to promote Tregs in the peripheral blood, lymph node, and spleen (Figures 17, 19, and 20). There was a corresponding increase in T cell exhaustion Figures 18, 21, and 22. CD20+ T cells were thoroughly depleted in all compartments by antibodies targeting CD20. They remained depleted even within the spleen two weeks after dosing in the high dose DFI205, and the combination Ofatumumab/DFI105 group, Figure 23. The combination treatment sustained the deepest depletion of these T cells.

**Figure 17:**
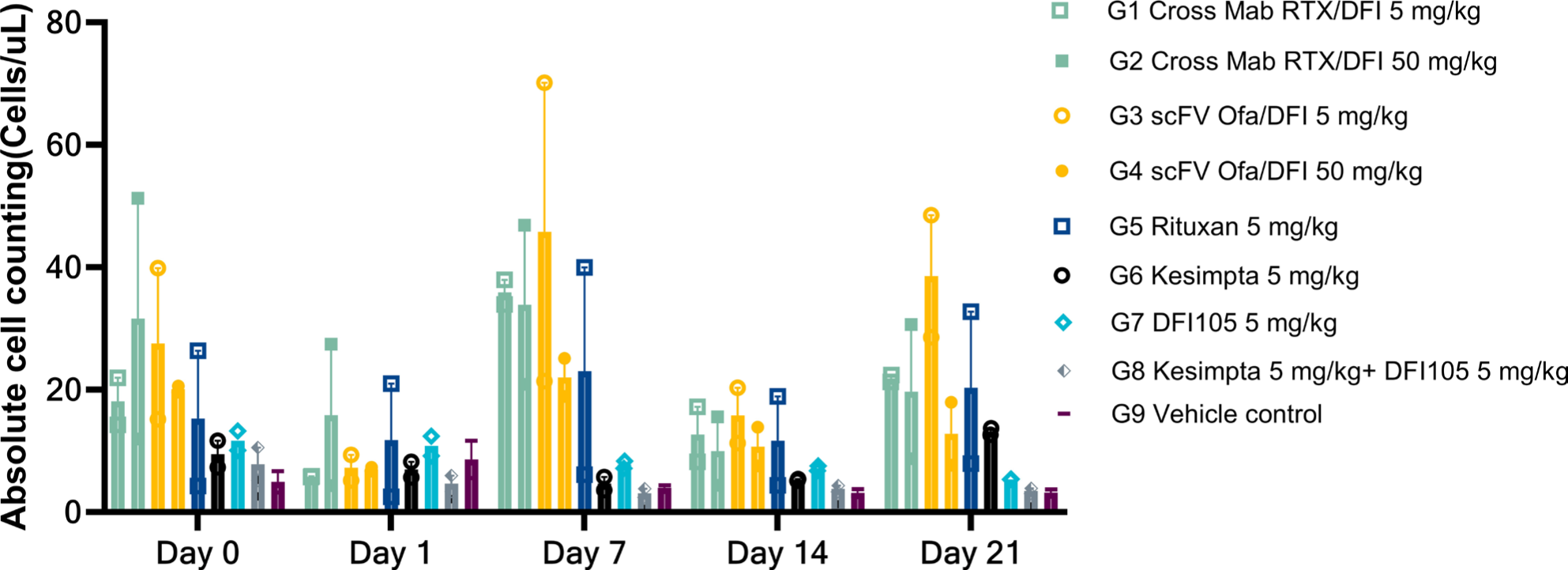
Blood Tregs During the Study.

**Figure 18:**
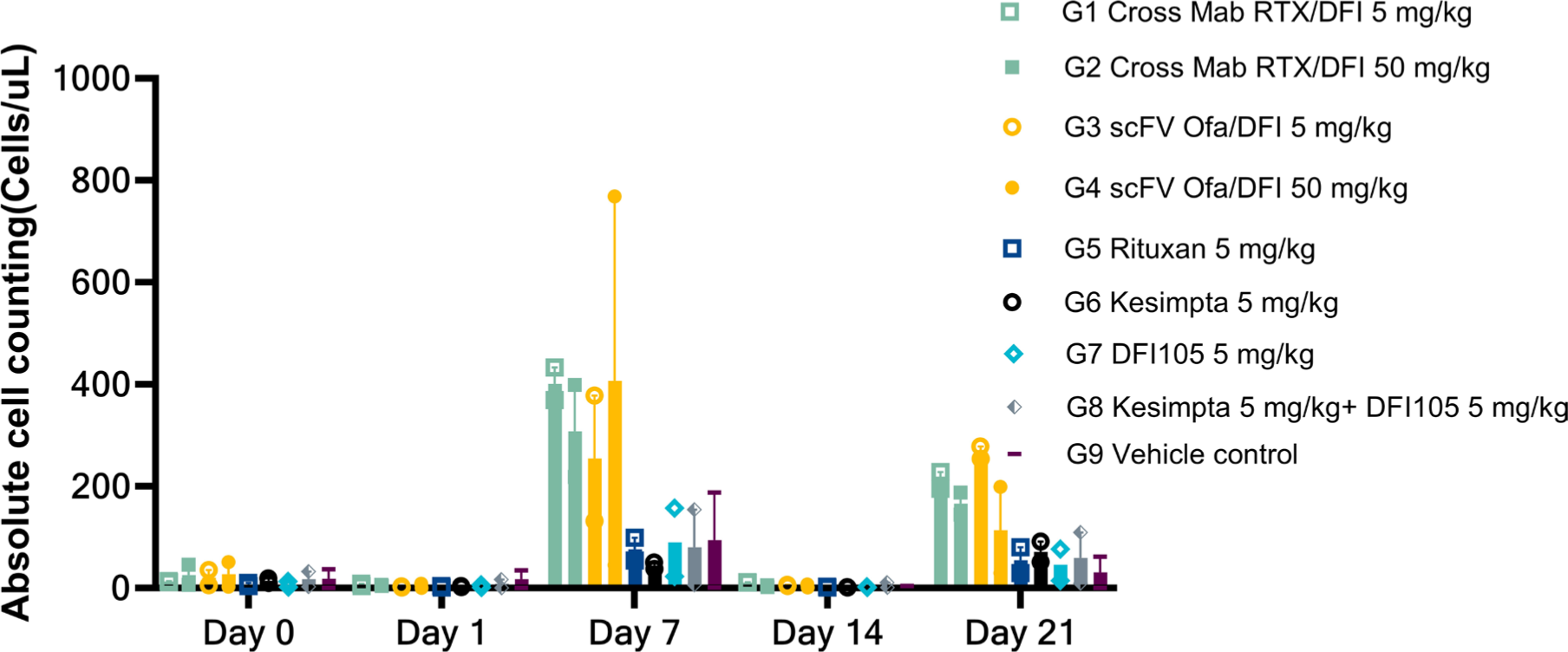
Exhausted T Cells during the NHP Study.

**Figure 19:**
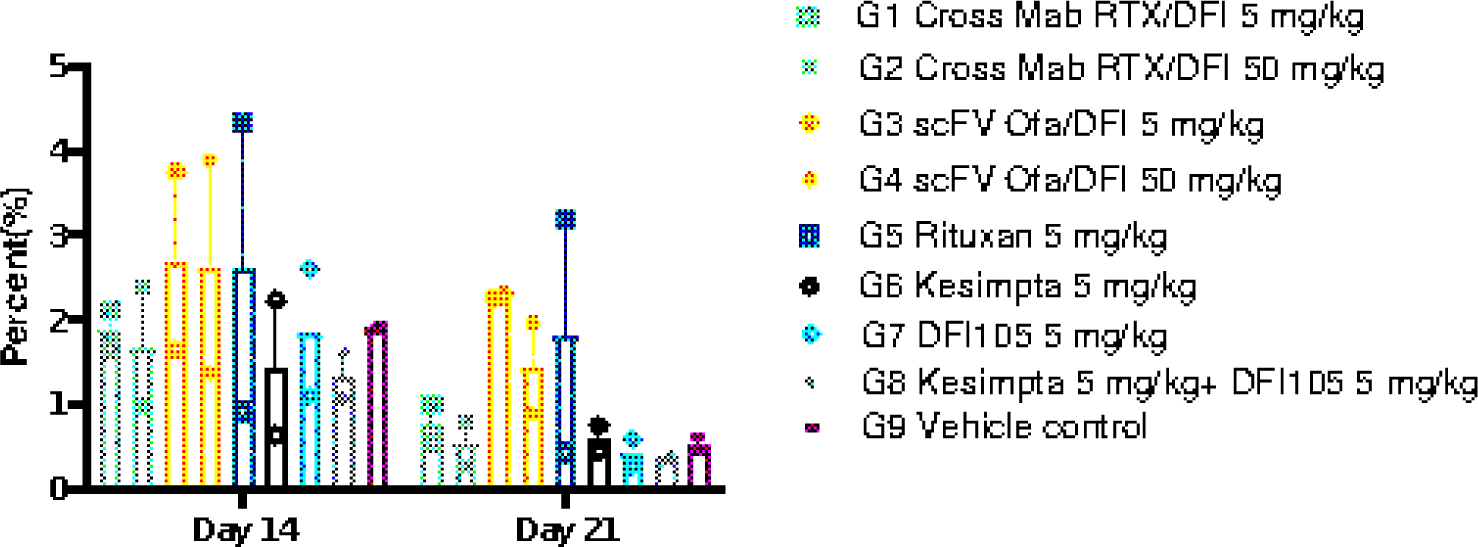
Lymph Node Regulatory T Cells One and Two Weeks after Dosing.

**Figure 20:**
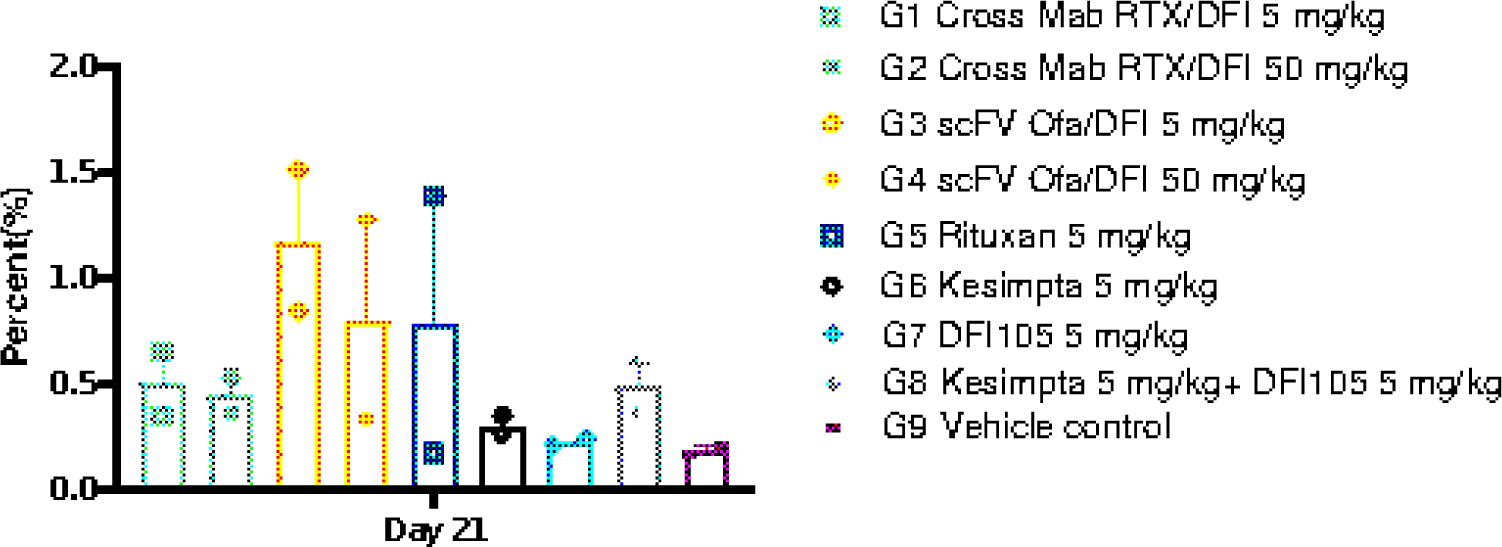
Treg Population in Spleen Two Weeks After Last Dose.

**Figure 21:**
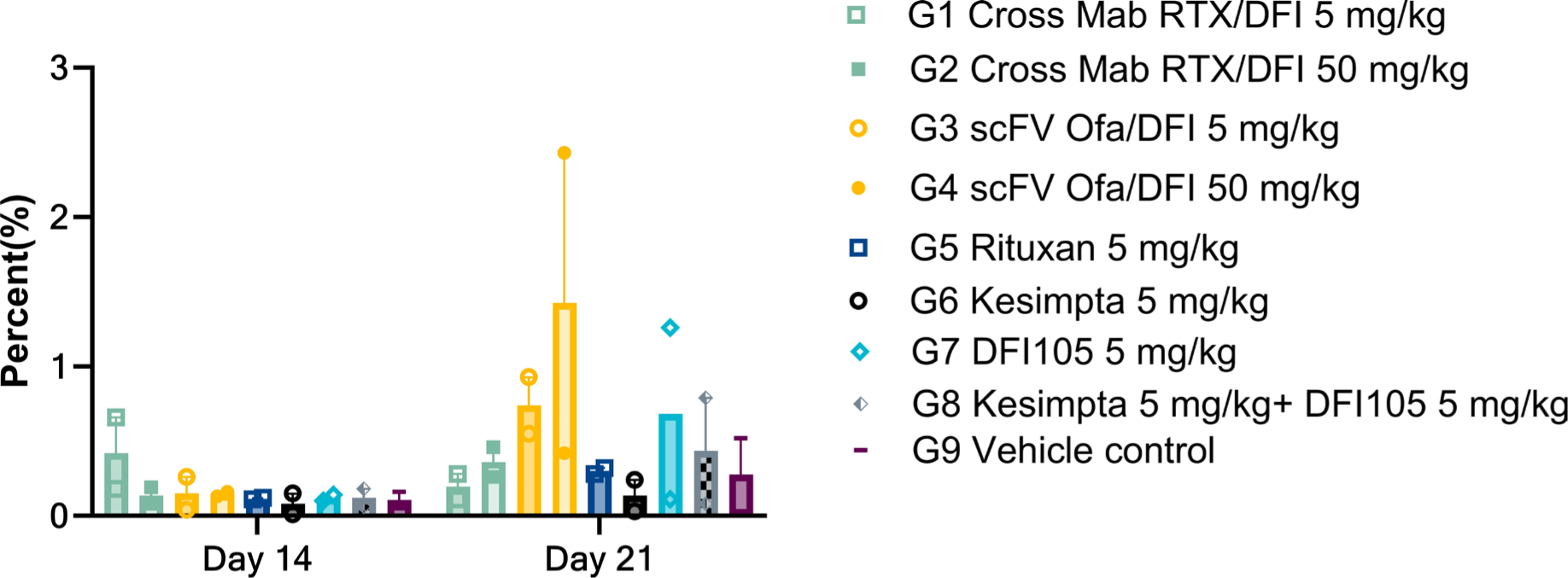
Exhausted T Cells Observed in Lymph Nodes.

**Figure 22:**
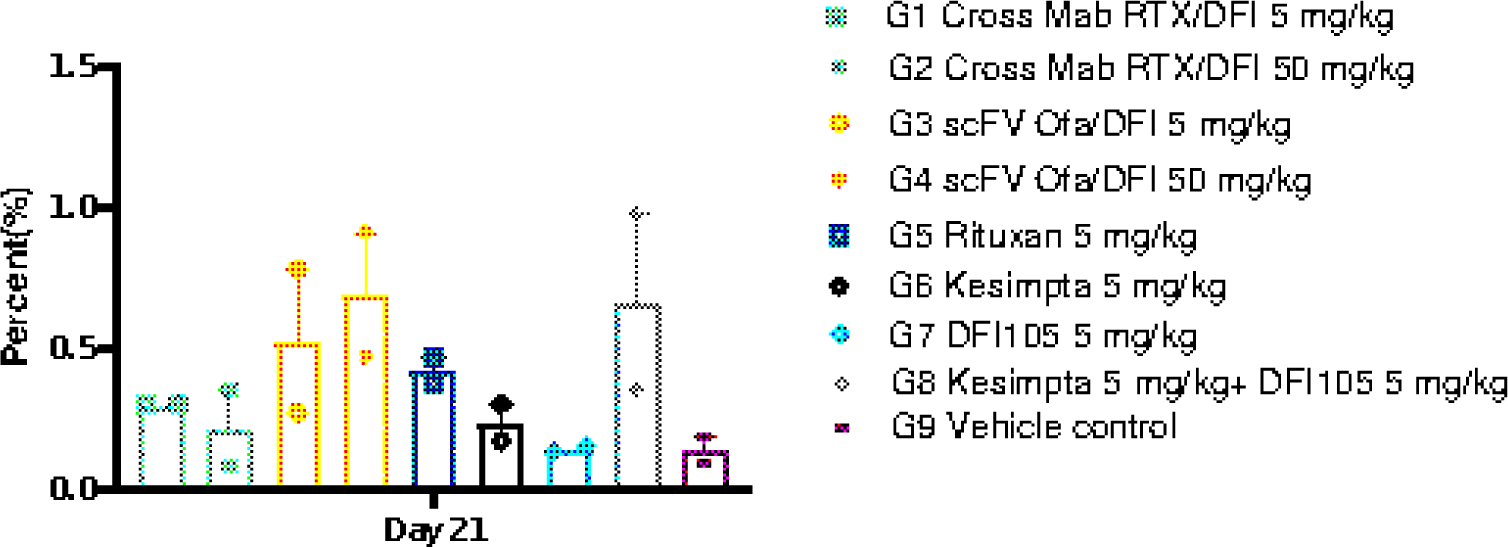
T Cell Exhaustion in the Spleen 2 Weeks after the final dose.

**Figure 23:**
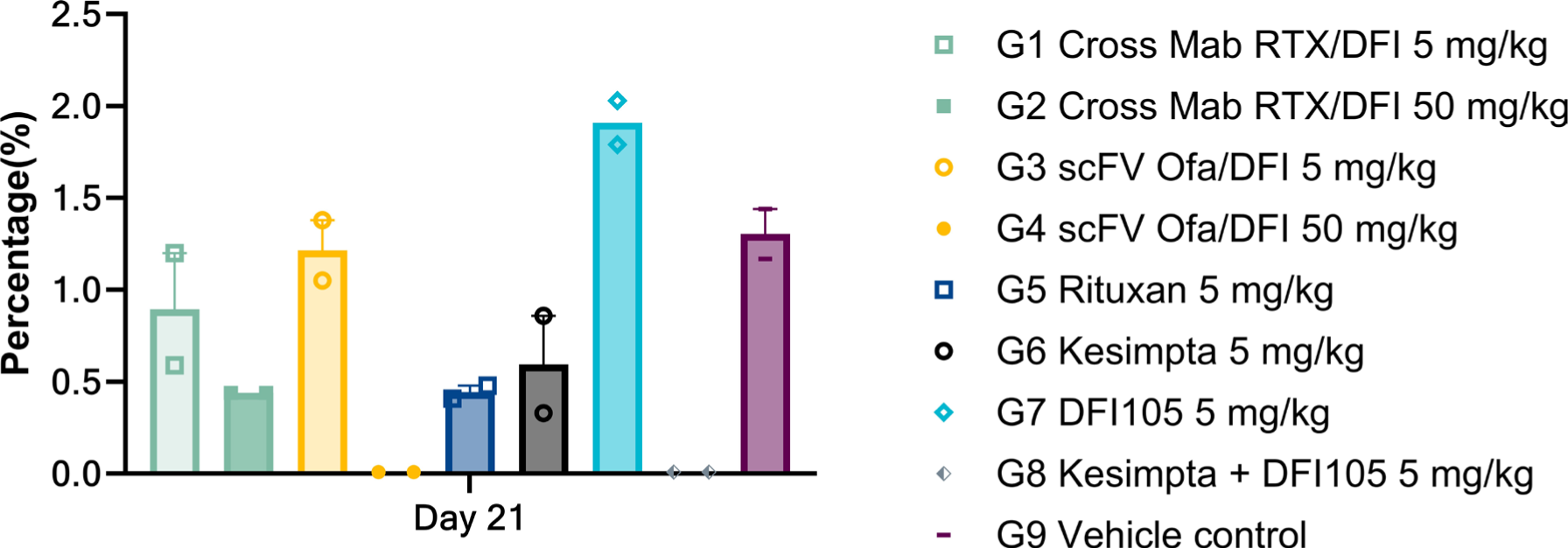
CD3+CD20+ T Cells in NHP Spleen At termination 2 weeks after the last dose.

### Immunohistochemistry

Immunohistochemistry (IHC) analysis was performed on lymph node and spleen tissues to characterize further immune cell distribution and tissue-specific responses to antibody treatments. Representative CD19+ and Ki67 stains from lymph node and spleen are provided in Figures 25 and 26. Results demonstrated distinct differences in cell populations, consistent with systemic flow cytometry findings. The 50 mg/kg dose of DFI205 and concurrent administration of 5 mg/kg doses of Ofatumumab and DFI105 demonstrated consistently superior B-cell depletion effects across both spleen and lymph nodes. Both groups had the lowest density of CD19+ cells per mm^2^ in lymph node and spleen IHC staining, Figure 24.

**Figure 24:**
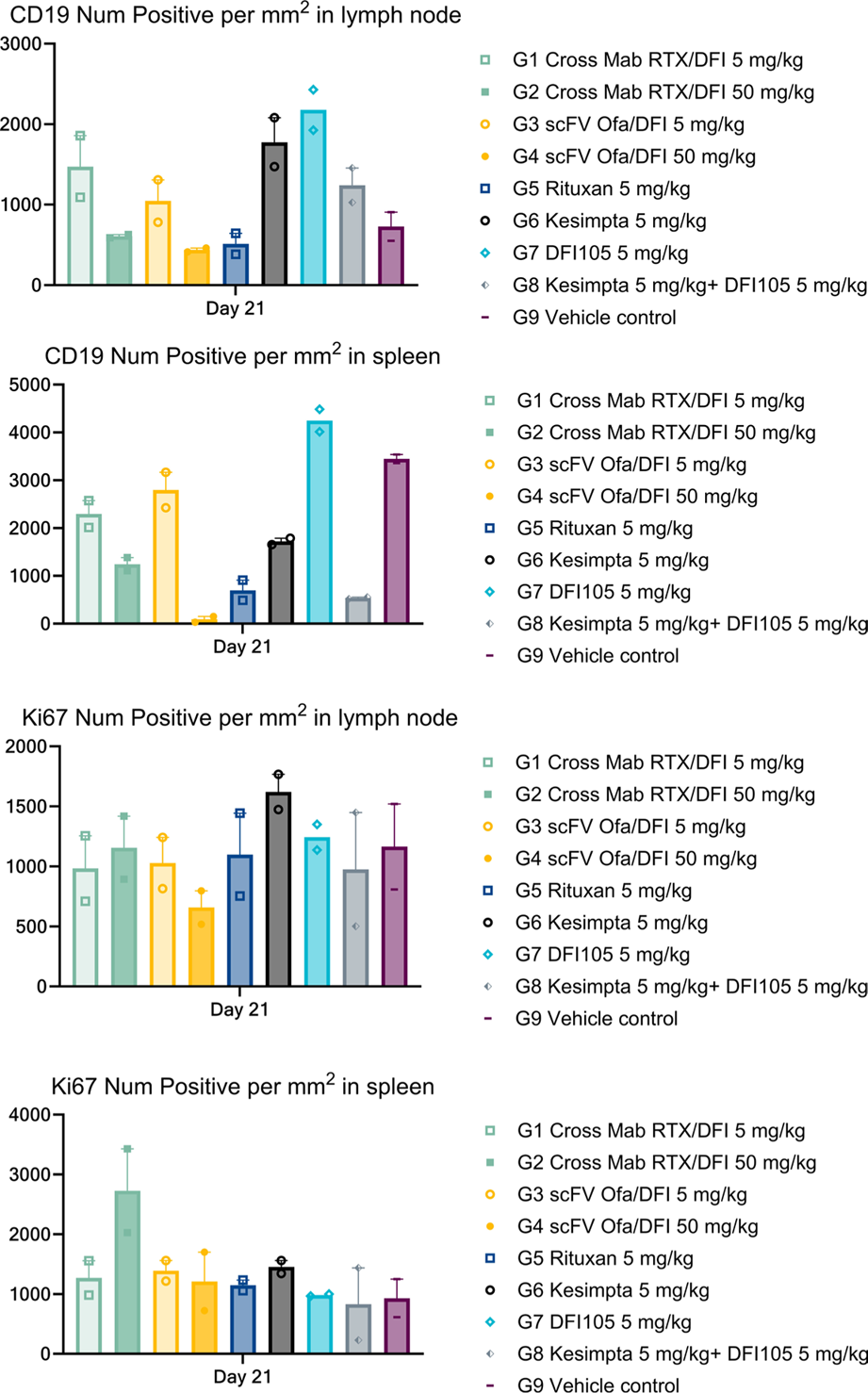
Quantitative Assessment of Tissue Resident CD19+ and Ki67+ in IHC Staining.

**Figure 25:**
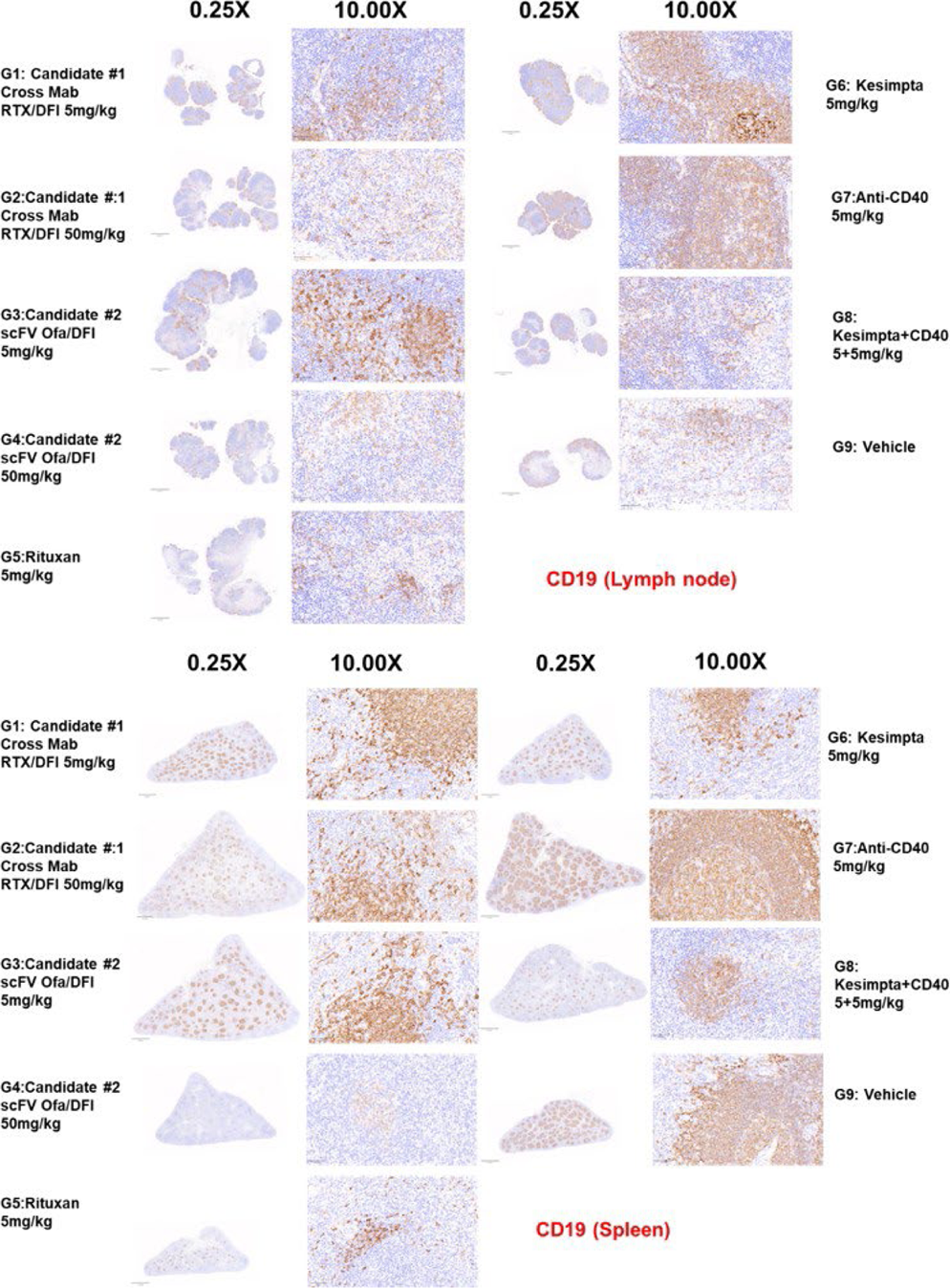
Representative pictures of lymph node and spleen CD 19 IHC staining.

**Figure 26:**
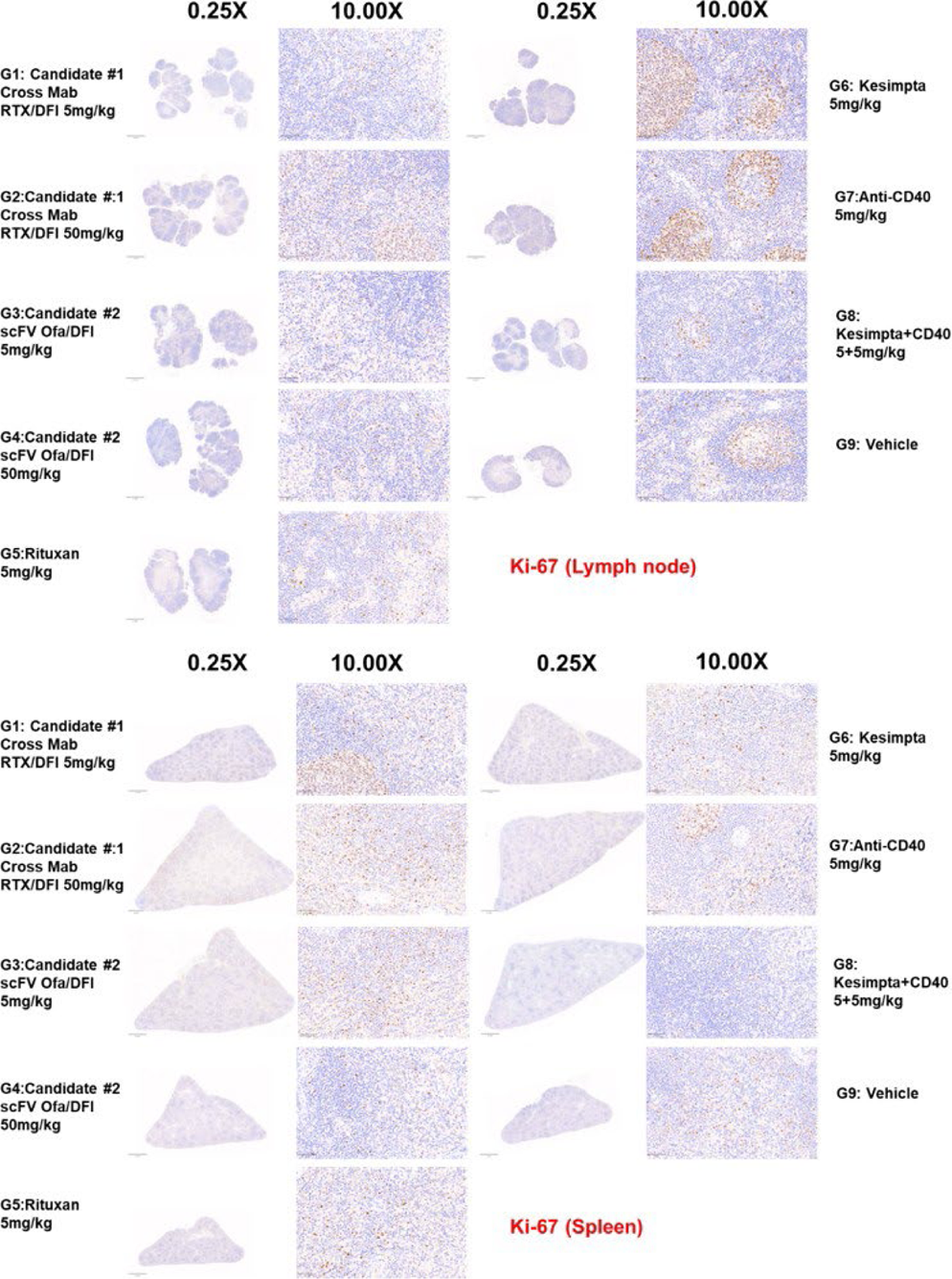
Representative pictures of lymph node and spleen KI-67 IHC staining.

#### Anti-Drug Antibody (ADA) Results

Across all groups, serum samples (Days 0, 7, 14, 21) were screened by a validated MSD bridging assay, and any putative positives were confirmed with an antigen-specific monkey IgG assay. In total, 19/72 samples exceeded the plate-specific screening cut-point, indicating putative ADA positivity; all of these were confirmed positive except for two samples collected on Day 7 from one DFI201 and one DFI205 treated animal. Results are summarized in Table 4.

**Table 4:**
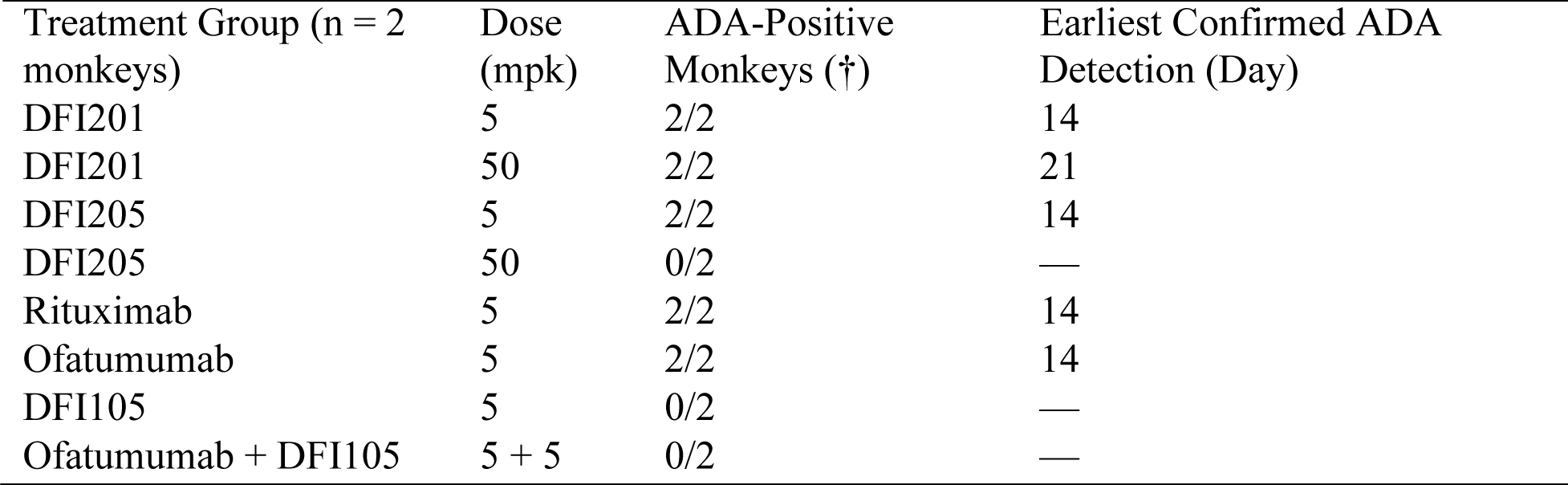
Summary of Anti Drug Antibody Occurrence.

The study drugs were human, or humanized IgGs. Monkeys developed ADA to DFI201 and DFI205 at the low dose, and Rituximab and Ofatumumab. ADA was absent in animals receiving high-dose DFI205 (50 mg/kg), anti-CD40 DFI105 alone, or the combination of Ofatumumab + DFI105. Confirmed ADA appeared between Day 14 and Day 21 and did not correlate with any adverse safety signals.

#### Safety and Tolerability

All treatment groups displayed acceptable safety profiles with no significant adverse events. Mild transient changes in hematology and clinical chemistry parameters resolved without intervention.

## Discussion

The treatment of monkeys with abti-CD20/CD40 bispecific and combined treatment with anti-CD20 and anti-CD40 antibodies was shown to be safe, without cytokine release syndrome, thrombotic events, or other negative safety signals.

The observation of anti-drug antibodies to anti-CD20 antibodies was consistent with testing human or humanized antibodies in non-human primates^30,31^. These do not correlate with the very low incidence clinical observation of ADAs in treatment with ofatumumab and rituximab^32,33^. ADAs were not observed in monkeys treated with anti-CD40 DFI205 alone or in combination with ofatumumab. ADA to anti-CD40 antibodies may occur when serum concentrations drop below the minimum effective dose level^34^. These antibodies, and the high dose of DFI205, induced sufficient immunosuppression to avoid ADAs in a set of monkeys that seemed prone to immunogenic responses. The bispecific half-lives were somewhat lower than the monospecific CD20 and CD40 antibodies, and showed clearance rates inversely proportional to dosage, suggesting target-mediated disposition as has been observed with ofatumumab^35,36^ and rituximab^37^.

The bispecific and combination treatment demonstrated full in vivo ability to execute the MOA of anti-CD20 monotherapy by providing rapid and sustained depletion of CD19+CD20+ in peripheral blood, with nadirs reached by Day 1 and minimal repopulation by Day 21. The bispecifics showed an uptick in B cell population comparable to CD20 mAbs, indicating that this depletion is reversible and safe. The clearance of peripheral and deep tissue B cells removes pathogenic and autoreactive B cells, effectively resetting the immune landscape, and treating a number of autoimmune diseases like multiple sclerosis^38,39^, rheumatoid arthritis^40^, and lupus^41^. Present treatments are not cures because tissue-resident and some memory B cells are out of the reach of conventional anti-CD20 antibody therapy^42–45^. Surviving B cells repopulate the peripheral immune system, and pathogenic memory B cells continue to present autoreactive antigens to T cells and perpetuate the disease state^46,47^.

Combining anti-CD20 and anti-CD40 provides superior depletion of memory B cells across blood, bone marrow, lymph nodes, and spleen, effectively eliminating CD27+ memory subsets. IHC quantification showed the lowest CD19+ B cell density in spleen and lymph nodes in high-dose DFI205 and combination groups, affirming deep tissue penetration and depletion. The bispecific’s efficacy relates in part to its dual-targeting nature and the broad lineage of B cells expressing CD20 and/or CD40 in relatively close physical proximity^24,48^. However, the success of the combination emphasizes some synergistic mechanisms from combining CD40 inhibition with CD20+ cell depletion. CD40 is expressed on endothelial cells and has functions related to cell migration from the skin to lymph nodes^49^ and angiogenesis^50,51^, in addition to pro-inflammatory signaling. Macrophages typically patrol deep tissues^52^. It is possible that while allosterically bound to CD40, mAb facilitated transport deep into tissues. The CD40 antibody may also engage monocytes and macrophages by crosslinking from CD40 to B Cells, and promote cell death^53,54^.

The bispecific antibodies demonstrated a unique ability to deplete non-conventional monocytes. In vivo, this manifested as a transitory reduction in the population of non-classical monocytes before homeostasis was established for the remainder of the study. Monocyte populations have short half-lives: a day for classical monocytes and a week for non-classical monocytes^55^. The continuous repopulation of these cells does not allow counts to serve as a surrogate for quality. Rheumatoid arthritis and other monocyte linked autoimmune disease may benefit from a partial reset of homeostaisis^56^. Concurrent depletion of deep tissue and bone marrow resident antigen-presenting cells may provide synergistic benefit.

The CD20*×*CD40 bispecific antibodies resulted in a pronounced modulation of dendritic cell (DC) populations, with a significant reduction in both myeloid DC1 and DC2 subsets in peripheral blood and lymphoid tissues by Day 21. This is likely a result of a dual mechanism where the anti-CD20 arm likely mediates direct cytotoxic depletion of CD20-expressing DC precursors^57^, while CD40 engagement appears to induce phenotypes that alter DC survival and function^58^. Ligation of CD40 on mature DC is known to enhance expression of costimulatory molecules. (CD80, CD86) and drive IL-12p70 secretion, thereby promoting T cell priming and regulatory T cell differentiation^58–60^. Inhibition of CD40-CD40L signaling prevents the creation of antigen-specific T helper cells^61^.

CD20*×*CD40 bispecific antibodies demonstrated a unique modulation of T-cell compartments, expanding regulatory T cells (Tregs) in both peripheral blood and lymphoid tissues with a 2-fold increase on Day 21 compared to baseline. Neither monotherapies nor the simple combination regimen provided the same Treg expansion. The frequencies of conventional effector CD4^+^ and CD8^+^ T cells remained essentially unchanged, indicating that the bispecifics expand Tregs without broad T-cells depletion. Regulatory T cells are central to maintaining self-tolerance and preventing autoimmune pathology^62,63^ deficiencies or functional impairments in Tregs have been causally linked to diseases such as type 1 diabetes^64^, multiple sclerosis^65^, and rheumatoid arthritis^66^. CD20*×*CD40 bispecifics offer a promising strategy to restore immune homeostasis in autoimmune disorders, potentially achieving durable disease control through rebalancing of regulatory and effector populations, rather than mere effector cell depletion. DFI205 showed a potential additional benefit by progressing antigen-presenting T cells to exhaustion, eliminating potential autoreactive T cells^67^.

All antibodies targeting CD20 demonstrated an ability to deplete CD3+CD20+ T cells, a small population of T cells implicated as pathogenic in MS and other autoimmune diseases^4,68^. The bispecific’s maintenance of this ability sustains its therapeutic potential. No direct measurement of pathogenic CD40+ T cell depletion was made^69,70^, but it follows that since the bispecifics

They are capable of depleting CD40-expressing cells and T cells; it is highly likely that they can address this population.

The multiple immunomodulatory mechanisms employed by the bispecifics demonstrated practical effect in a T cell–dependent antigen challenge using keyhole limpet hemocyanin (KLH). Both CD20*×*CD40 bispecific antibodies profoundly suppressed primary humoral responses: animals treated with DFI201 or DFI205 mean anti-KLH IgG titers of <5% relative to vehicle controls by Day 21, whereas rituximab and ofatumumab monotherapies only achieved ∼40–60% reductions in titer relative to vehicle. Anti-CD40 monotherapy DFI105 provided immunosuppression higher than an equivalent dose of BsAb. The near-complete abrogation of KLH-specific antibody generation underscores the dual capability of CD20*×*CD40 bispecifics: not depletion of B cells but also CD40 inhibition of adaptive humoral immunity. This blockade of T cell–dependent antibody formation suggests a powerful mechanism for re-establishing tolerance in autoantibody-driven diseases.

## Conclusion

Our study demonstrates that bispecific antibodies integrating CD20-mediated depletion with antagonistic CD40 inhibition synergistically reshape both humoral and cellular immune compartments. By achieving profound and sustained clearance of B cell and dendritic cell populations, transient resetting of monocyte homeostasis, and expansion of Tregs without overt safety concerns. CD20*×*CD40 bispecific antibodies provide a versatile platform for inducing antigen-specific tolerance. These findings support further development of this approach for the treatment of autoimmune disorders and allograft rejection, where coordinated targeting of multiple immune lineages may overcome limitations of current monospecific therapies.

## Notes

### Competing Interest Statement

The authors have declared no competing interest.

